# Time-resolved inference of gene regulatory networks underlying human cranial neural crest development suggests novel risk genes for orofacial clefting

**DOI:** 10.64898/2026.06.25.734423

**Authors:** Michael Eibl, Susanne Theiss, Hjörleifur Einarsson, Christian Skov Vaagensø, Robert Krautz, Maren Gehringer, Anna Siewert, Youcheng Zhang, Álvaro Rada-Iglesias, Julio Saez-Rodriguez, Carl Herrmann, Kerstin U. Ludwig, Robin Andersson, Magdalena Laugsch

## Abstract

Cranial neural crest cells (CNCCs) play a central role in shaping the human head and face. Aberrant CNCC differentiation contributes to craniofacial birth defects, particularly non-syndromic cleft lip with or without cleft palate (nsCL/P), one of the most common congenital disorders. Although the number of genetic variants associated with this condition is steadily increasing, it remains challenging to determine if and how these variants may contribute to disease development. The majority of these variants lie within non-coding regulatory elements that govern cell-type and stage-specific gene expression, which is orchestrated by dynamic gene regulatory networks (GRNs). Despite extensive work in model organisms, a time-resolved, multi-omics perspective of GRNs controlling CNCC differentiation in a human system is still lacking.

To fill this gap, we generated paired transcriptomic and chromatin accessibility data at four timepoints during *in vitro* differentiation of CNCCs derived from human induced pluripotent stem cells. Integrating these two modalities enabled time-resolved inference of GRNs and identification of dynamic regulatory relationships, including stage-specific roles of core transcription factors. Leveraging these time-resolved GRNs, we mapped 29 nsCL/P associated variants linked to 70 putative target genes, with 40 located outside the associated genomic loci, suggesting novel distal regulatory relationships. Integration of these data with complementary time-course scRNA-seq data revealed an ectomesenchymal-biased subpopulation of CNCCs as particularly sensitive to genetic variants associated with nsCL/P.

We provide a time-resolved inference of GRN in human CNCC differentiation, allowing us to determine the dynamics of stage-specific core regulatory programs that are otherwise missed in analyses based on a single time snapshot. To our knowledge, the data represent the first multi-omics map of human CNCC with temporal resolution, which expands the understanding of early human craniofacial development, refines variant-to-gene assignment, prioritizes candidate risk genes and cell states relevant to nsCL/P. Our findings demonstrate the relevance of studying the dynamics upon differentiation rather than just one fixed timepoint and offer a valuable basis for further investigation of non-coding variation in CNCC-related disorders.

## Introduction

The neural crest (NC) is a transient, multipotent cell population unique to vertebrate embryonic development. Originating at the border of the neural plate and non-neural ectoderm during neurulation, NC cells undergo induction, specification, and an epithelial-to-mesenchymal transition (EMT). These processes enable NC cells to delaminate from the dorsal neural tube and migrate throughout the embryo in stereotypical patterns. Ultimately they differentiate into a wide range of cell types, including cartilage, bone, peripheral neurons and glia, pigment cells and endocrine tissues [1–3]. One of the key aspects of NC development, extensively studied in model organisms such as frog [4], chicken [5] or zebrafish [6], is tightly controlled spatiotemporal gene expression governed by gene regulatory networks (GRNs). Species-specific differences, including enrichment of core transcription factors such as *Nr2f1* and *Sox9* in mouse but not chick pharyngeal arch NC cells [7], underscoring the need for studies of species-specific GRNs, particularly in the context of human disease. While first studies have leveraged data from human embryos to characterize gene expression dynamics in craniofacial morphogenesis [8], the concomitant profiling of chromatin accessibility data covering a tight window of early neural crest development in a human system is lacking.

NC evolution, particularly in the cephalic region giving rise to cranial NC cells (CNCCs), has been driven by the need for specialized craniofacial structures, such as a complex jaw, that equipped animals for a predatory lifestyle [9], giving rise to the majority of the craniofacial mesenchyme. CNCCs contribute to the skull vault, upper and lower jaw (maxilla and mandible) as well as the frontonasal prominence, the inner ear or the hyoid bone [10]. Humans have particularly high intra-species craniofacial variability, with *Hominoids* displaying highest variance in cranial shape even among primates [11]. This has been suggested to have evolved to serve social purposes [12] including a self-domestication process, with specific regulators like the chromatin remodeller BAZ1B playing a central role [13]. Craniofacial morphology is a highly polygenic trait and genome-wide association studies (GWAS) have revealed a breadth of genomic variants associated with the shape of face and cranium [14–18]. Several face shape variants were annotated within their gene regulatory context, showing the relevance of TCF or ALX family of transcription factors (TFs) in craniofacial morphogenesis in a human model system [19]. In addition, studies of individual TFs, including SOX9, NR2F1/2 and TFAP2A demonstrated their essential contribution to human CNCC formation in health and disease [20–23]. Hence, establishing functional links between genomic variants and craniofacial morphology is crucial to understand not only general human face variation but also the etiology of craniofacial conditions such as craniosynostosis, frontonasal dysplasia or orofacial clefting [24]. Indeed, the genetic basis for philtrum width in faces is significantly overlapping with the genetic basis for non-syndromic cleft lip with or without cleft palate (nsCL/P) [25]. NsCL/P is among the most prevalent congenital birth defects, affecting around 1/1000 live births [26] and leads to feeding difficulties, negatively affects speech development and overall quality of life [27], typically requiring early surgical interventions. While new GWAS data continuously expand the list of variants associated with nsCL/P [28–31], how most of these variants functionally contribute to the etiology of the condition is not fully determined [29]. This functional gap, in addition to ethical constrains limit the study of cell type and developmental timing of nsCL/P determination in the human embryo, which may differ between variants, making human *in vitro* models indispensable.

Most of the associated variants identified by GWAS on facial traits or diseases are located in the non-coding genome, presumably in regulatory elements (REs), that are bound by TFs to control expression of specific target genes (TGs) [32]. Notably, TFs may act in a highly synergistic manner even within the same RE [33], underscoring the need for a global gene regulatory perspective beyond occupancy profiling of individual TFs. GRN-based links of genomic variants to TGs have recently been used to functionally annotate single nucleotide polymorphisms (SNPs) associated with normal human face variation and craniofacial dysmorphology [19,34]. These studies were, however, limited to a single time point in CNCC development, corresponding to already strongly ectomesenchymally-biased CNCCs or were performed in the zebrafish periderm – a non-neural crest contributor to orofacial morphogenesis.

To fill this gap, we inferred time-resolving GRNs governing multiple stages of human CNCC development, particularly induction in the neuroepithelium, specification, EMT and migration, and after acquiring their cranial identity. For this purpose, we integrated paired transcriptomics and chromatin accessibility data during human CNCC differentiation from human induced pluripotent stem cells (hiPSCs). We employed cap analysis of gene expression (CAGE), which detects 5’ capped RNA and maps transcription start sites (TSSs) [35]. Importantly, CAGE also captures non-polyadenylated transcripts such as enhancer RNAs (eRNAs) at active enhancers, thereby providing an orthogonal layer for RE identification by assessing enhancer activity in addition to chromatin accessibility measured by Assay for Transposase-accessible chromatin (ATAC) sequencing [32,36,37]. We validated inferred TGs for core human CNCC regulators NR2F1 and TFAP2A within the obtained GRNs using published ChIP-seq data and inferred core TFs at each stage in an unbiased manner by non-negative matrix factorization (NMF). Subsequent mapping nsCL/P-associated non-coding variants onto the GRNs revealed both known variant-target gene links and novel putative links to target genes that have been previously linked to craniofacial dysmorphology, but not to these susceptibility regions. Moreover, integrating our GRNs with time-course single-cell RNA-sequencing (scRNA-seq) data and nsCL/P-associated variants enabled prediction of both known and novel nsCL/P risk genes and highlighted hCNCC biased towards ectomesenchymal fates as particularly sensitive to genetic variants associated with nsCL/P. Our findings underscore the value of GRNs inferred from time-resolved multi-omics data in a human-specific system to deepen our understanding of the genetic basis of NC-associated diseases, i.e. neurocristopathies, in general.

## Results

### hiPSC-derived human CNCCs recapitulate major features of NC development

Although several protocols for *in vitro* differentiation of hiPSC into human CNCC have been reported providing valuable insights into human CNCC development, most omit the EMT [38–43], a hallmark of the NC biology. Therefore, we employed a protocol that faithfully recapitulates EMT, enabling the study of early stages of human CNCC development including induction, specification, and migration [20–23,29,44,45]. We systematically dissected the temporal dynamics of gene regulation during human CNCC differentiation by generating transcriptomic and chromatin accessibility data at four selected timepoints (d5, d8, d11 and d18, hereafter referred to as p2). These timepoints encompass the presence of already formed floating neuroectodermal spheres (NEC), enabling ongoing NC induction and specification (d5-8). EMT and delamination occur upon NEC attachment to the culture dish (d5-11), followed by NC outgrowth and migration (d8-11). Finally, the adherent and migrating NC cells acquire cranial identity by the second passage (p2) (Supplementary Figure 1a) [20–23,44,45]. Paired CAGE and ATAC sequencing data were obtained in triplicates at each timepoint to capture transcriptional and chromatin accessibility dynamics throughout human CNCC development (Supplementary Figure 1a). As for CAGE, the highest transcription start site (TSS) signal was observed at annotated promoter regions, while eRNAs were detected in both intergenic and intronic regions. The average CAGE signal intensity at promoters was approximately 1.3-fold greater than in other regions (Supplementary Figure 2). The ∼300,000 ATAC peaks per sample showed similar distribution around annotated TSSs and across genomic features (Supplementary Figure 3). In accordance with previous work [46], the majority of ATAC peaks mapped to promoter and intergenic regions.

The differentiation timepoints constitute the major source of variability in both gene and eRNA expression as well as chromatin accessibility (Figure 1a-b, Supplementary Figure 3a-c) reflecting the nature of the human CNCC differentiation protocol. At d5, all floating NECs are already formed. From this timepoint on, the NECs spontaneously begin to attach to the culture dish to give rise to the first migratory NC cells. By d8, the cell population is mixed, comprising immature and more mature outgrowing NC cells alongside differentiating NECs that intermittently attach and detach, giving rise to new NC cells. These dynamics are clearly observed in the chromatin accessibility data (Figure 1b), and less pronounced in the beginning of the differentiation in the transcriptomics data (Figure 1a), consistent with chromatin accessibility changes preceding gene expression changes [47]. At day 11, most of the mature NECs have attached and flattened, while the majority of NC cells have delaminated and migrated away from the NECs. By p2, cells are more homogeneous, consisting of human CNCCs as they were previously passaged at d11 and a few days later to fully eliminate NECs. These dynamics are further proven by dynamically expressed marker genes (Supplementary Figure 1b). The transcriptional transition from neuroepithelial to NC states is reflected in a progressive decrease in key neuroectodermal markers such as *SOX2*, *PAX6*, *SIX3*, and *CDH2* (see neural rosettes in Supplementary Figure 1c) and simultaneous induction (*PAX7*) and specification (*SOX10*) of NC between d5-11 (Supplementary Figure 1c-d). The upregulation of *SOX9*/*TFAP2A*/*TWIST1*/*NR2F1* together with increased expression of migratory markers (*RHOB*/*SNAI2*/*ZEB1*) show the acquisition of human CNCC identity by p2 (Supplementary Figure 1b) [2]. Moreover, the transcriptional dynamics align with the regionalized induction of a SOX10 and TFAP2A-positive population within d8 NECs and heterogeneous SOX10 protein levels by d11 as confirmed by co-immunofluorescence (Supplementary Figure 1c-d). These data therefore provide a unique and valuable basis to elucidate GRNs in a human-specific system using human CNCCs derived from hiPSCs.

**Figure 1.**
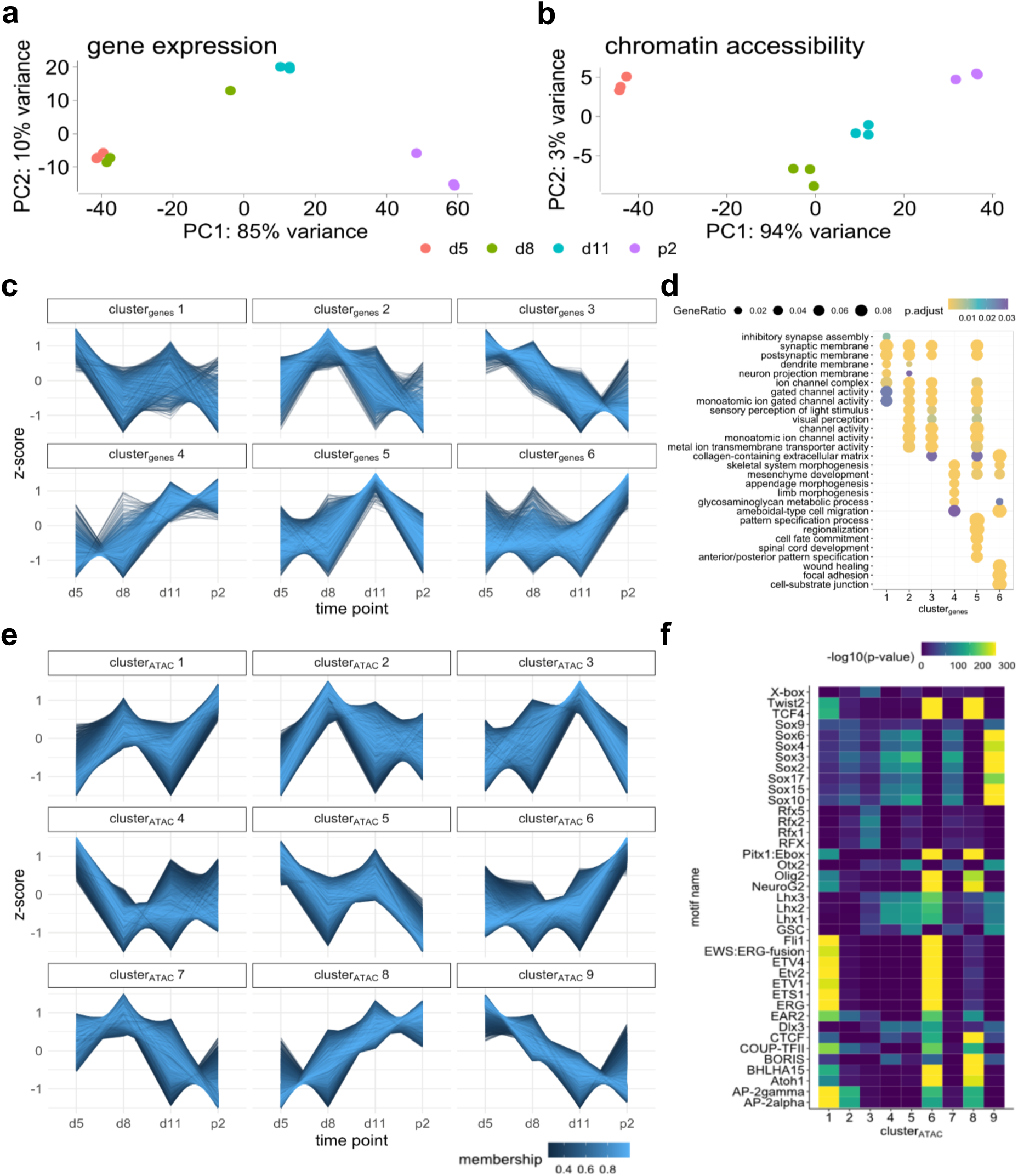
Gene expression and chromatin accessibility dynamics at d5, d8, d11 and p2 during hiPSC-derived human CNCC differentiation. **a-b** Principal component analysis (PCA) of the CAGE **(a)**, and ATAC-seq data **(b)** indicates a continuous differentiation trajectory corresponding to the timepoints. **c** Dynamically expressed genes (log2FC > 1 when comparing one timepoint to any other) were clustered together according to similar temporal expression profiles by fuzzy c-means clustering. Relative expression per gene across the four time points is shown as z-scores. **d** Gene ontology analysis was performed per cluster. **e** Dynamically accessible regions (log2FC > 1 when comparing one time point to any other) were similarly clustered according to their temporal accessibility profiles using fuzzy c-means clustering. Relative accessibility values per region across the four time points are shown as z-scores. **f** Cluster-wise TF motif enrichment analysis performed using HOMER indicates putatively regulating transcriptional regulators specific to regions with distinct accessibility dynamics.

### Mapping of gene expression and chromatin accessibility dynamics during development of human CNCCs

To generate a time-resolved map of expression and accessibility profiles during human CNCC differentiation, we first employed a soft-clustering approach (fuzzy c-means clustering) on our CAGE and ATAC-seq data (Figure 1c-f). In the transcriptomics data, we found six clusters that capture the major patterns in gene expression dynamics. Clusters 4 and 6 contain genes associated with mesenchyme development and skeletal system morphogenesis, which are upregulated during the differentiation (Figure 1c-d). These include e.g., the collagen *COL1A1* and the ectomesenchymal regulator *FOXC2* (Supplementary Figure 5), consistent with the known ectomesenchymal characteristics and differentiation potential of CNCCs [1,2,48]. Cluster 1 and 3 include neuronal lineage associated genes transiently expressed only early in the differentiation, peaking at d5. These clusters highly express *CDH2* and low levels of synaptic genes (*GABRB2*, *GRIA1*, *SHANK2*) (Supplementary Figure 5), suggesting a mixed progenitor state with partial neural commitment, highlighting the heterogeneity of NECs. Cluster 2 genes with peak expression on d8 are enriched for genes associated with visual perception, including *EYA4* or *CRYBA1* (Supplementary Figure 5). Thus, our data strongly suggest that the neuroectoderm may have partially differentiated towards cells reminiscent of the optic cup *in vivo*, a process that notably relies on the cross talk between neural progenitors (NPC) and NC [49], highlighting the self-organizing principles governing the differentiation. In cluster 5, we identified patterning-associated genes with peak expression on d11 (Figure 1c-d, Supplementary Figure 1). These include low levels of *HOXA2* and *HOXA3* (Supplementary Figure 5), which pattern the cranial NC along the anterior-posterior axis. These genes are typically only expressed in pharyngeal arches (PA) 2 and 3, whereas PA1 CNCC and the mesencephalic stream of CNCCs are HOX-negative [50]. Overall, our data confirm the expression of predominantly anterior cephalic genes and highlight several features of self-organized diversification of the anterior neuroectoderm from which ultimately human CNCCs arise by p2, overall recapitulating important events *in vivo*.

Clustering of our ATAC-seq data revealed nine clusters of distinct accessibility dynamics (Figure 1e), which was followed by cluster-wise TF motif enrichment analysis using HOMER [51] (Figure 1f). Clusters 1, 6, and 8 contain putative human CNCC REs, enriched in motifs for the AP2, NR2 and TWIST family. Of note, the ETS superfamily, found in cluster 1 includes ETS1, ETV1/4, which have been strongly implicated in NC development [52]. We also observed strong enrichment of CTCF and BORIS (CTCFL) motifs in cluster 8, which exhibits the most linear increase in accessibility during the differentiation, suggesting that these factors play a key role in progressively rewiring chromatin to enable activation of differentiation-specific genes. Clusters 4, 5 and 9, on the other hand, contain regions with decreasing accessibility during differentiation and are enriched for motifs of NPC regulators of the SOX and LHX family of TFs, as well as OTX2. In cluster 3, we identified most accessible regions on d11, which are enriched for RFX family motifs (Figure 1f). This pattern aligns with the idea that d11 early migratory NC cells are likely to strongly induce ciliogenesis [53], whereas p2 cells may predominantly rely on ameboid-like crawling for locomotion (Figure 1c-d, cluster 6) consistent with previous studies from different model organisms [54]. Altogether, based on our ATAC-seq data analysis, we identified several well-known regulators of neuroectodermal and CNCC fates, highlighting strong conservation of these processes between species, in addition to poorly characterized regulators, which may have time-point specific roles, such as RFX family TFs.

In accordance with previously reported studies, we assume that the observed gene expression changes during human CNCC differentiation (Figure 1c-d) are partially driven by tissue- and timepoint specific enhancers [46]. Chromatin accessibility status alone (Figure 1e-f) is, however, not sufficient to determine enhancer activity [32]. Therefore, we quantified CAGE signal in promoter-distal accessible regions (5000 bp from annotated TSSs). Of these 19,281 distal REs, 6,174 showed CAGE signal in at least two time points, indicating their activity as enhancers. The average CAGE dynamics closely mirrored the average accessibility dynamics across the 9 clusters (Supplementary Figure 6a-b), in line with the close link between chromatin accessibility and enhancer expression and thereby activity. GREAT analysis revealed that these putative enhancers are located near genes governing differentiation and signal transduction (Supplementary Figure 6c), while HOMER analysis showed enrichment of similar motifs as for the global ATAC dynamics (Supplementary Figure 6d). The data highlight similar timepoint-dependent TF usage in both promoter-proximal and distal regions with and without CAGE signal as exemplified for two CNCC differentiation-associated putative enhancers (cluster 6) located at the *DLX1* locus, a gene consistently upregulated throughout the differentiation (Supplementary Figure 7b) and essential for the patterning of the jaw [55]. We observe little accessibility changes at the promoter, but instead at putative intronic REs (Supplementary Figure 6a). As shown in Supplementary Figure 7c, two of these overlap bidirectional CAGE-tags (eRNA expression). Putative REs with matching accessibility, eRNA and gene expression changes throughout the differentiation are therefore potential candidates for future locus-specific investigations. However, to determine global GRNs underlying the differentiation process we focused on all ATAC-seq identified putative REs, some of which may be missed by CAGE, as reported for lowly expressed eRNAs [56].

### Inference of GRNs and core TFs governing human CNCC differentiation

Motif enrichment analysis alone cannot reliably distinguish TFs of the same family (Figure 1e-f) and thus oversimplifies the regulatory complexity. To overcome this limitation, we integrated both TF and target gene (TG) expression data with the paired chromatin accessibility profiles using PECA2 to infer GRNs. Visualising TF-TG subnetworks per timepoint (Figure 2a-d), we focused on five master regulators per time point with well-known relevance for the neuroectoderm [57], the NC and the ectomesenchyme [2,19]. For clarity, only the top three TGs per TF with the highest trans-regulation scores (TRS) and the regulatory interaction strength between these core TFs are shown (Figure 2a-d). We observed strong combinatorial TF regulation, a general feature that ensures timepoint- and tissue-specific gene expression [58]. Consistent with positive autoregulation as a crucial feature during development that supports robustness of TF levels [59], we captured autoregulation of specific TFs, such as SOX2 on d5, SOX10 on d8, NR2F1 on d11 or TFAP2A in p2 (Figure 2a-d), coinciding with their expression peaks (Supplementary Figure 1b). SOX2 regulates several other neuroectodermal TFs, including ZIC2 and OTX2, while receiving input from ZIC2 and itself (Figure 2a). At d8, the anterior neuroectoderm regulator RAX is upstream of NC inducer PAX7 and NC master regulator SOX10 (Figure 2b). Early migratory NC cells at d11 (Figure 2c) show substantial cross-regulation between NC regulators NR2F1, cranial ALX4, and the mesenchymal regulator PRRX2 which may be essential for neuroectoderm and NC differentiation potential. Interestingly, HOXA2 is upstream of NR2F1 and PRRX2, indicating anterior positional regulation of CNCC identity [50]. Among the TGs strongly controlled by the known core TFs in p2 (Figure 2d), several other TFs were found, including the mesenchymal/EMT regulator PRRX1 [60] and PKNOX2, implicated in cochlear development [61] and expressed in melanocytes [62], suggesting a broader role of PKNOX2 in human CNCC development. The osteogenesis regulator *SIX1* is tightly controlled by core TFs in p2, consistent with the ectomesenchymal character of these cells and mutations in *SIX1* associated with craniofacial defects, including in Branchio-oto-renal syndrome [63]. LMX1B, important for periocular mesenchyme formation [64], is also strongly regulated by core CNCC TFs. Among non-TF TGs, the intermediate filament Nestin (NES), a marker of NPCs and NC stem cells [65], and stemness-associated MEX3A, expressed in proliferative CNCC [66] show strong regulation, indicating that this tight control may be essential for neuroectoderm and NC differentiation potential (Figure 2b-d). Overall, TGs with highest TRS per core TF are known key tissue regulators, suggesting that TRS correlates with developmental and functional relevance (Figure 2a-d).

**Figure 2.**
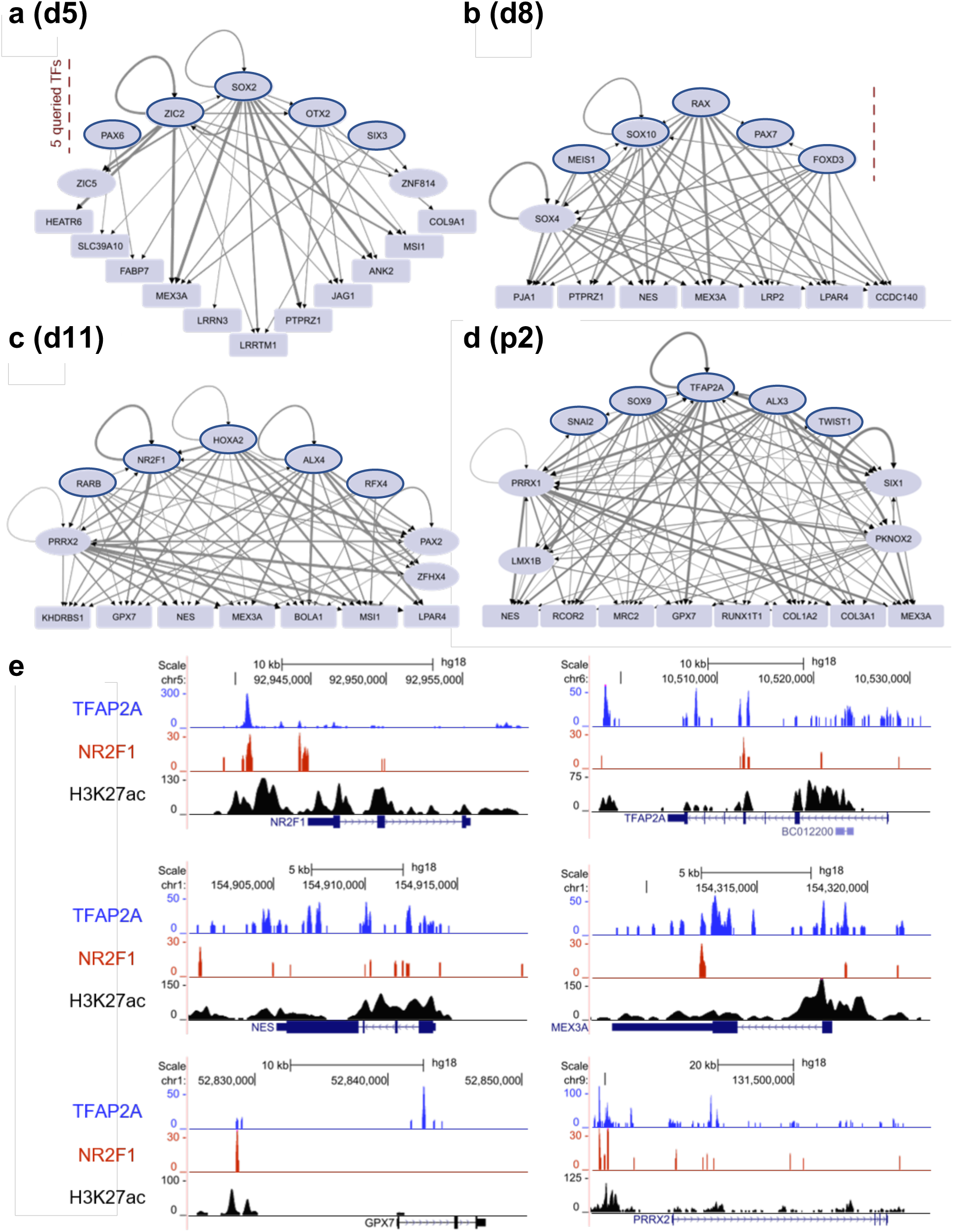
Gene regulatory network inference by PECA2 infers regulatory relationships and TF hierarchies in human CNCC development. **a-d** Visualization of selected inferred subnetworks per timepoint. Three TGs with the highest TRS were queried for a set of five TFs (encircled ellipses) with known relevance during human CNCC differentiation. Rounded rectangles indicate non-TF TGs. A cutoff of TRS=30 was applied for the TF-TG pairs. **e** NR2F1, TFAP2A, and H3K27ac ChIP-seq tracks from NC derived from human embryonic stem cells (hESCs) using the same protocol as ours [20] validate our inferred TGs of high TRS for d11 NC and TF binding at RE with active chromatin marks.

Feng et al. (2021) used published data from human CNCC in passage 4 (p4) to infer hReg-CNCC and validated their predictions with corresponding ChIP-seq data [19,23]. However, p4 exhibits a greater ectomesenchymal bias compared to the earlier timepoints, which we examined here. Hence, we probed the validity of our TF-TG pairs with high TRS for d11 (Figure 2a-d), matching available ChIP-seq data for NR2F1, TFAP2A, and the active chromatin mark H3K27ac [20]. As shown in Figure 2e, we confirmed virtually all inferred predictions based on TF binding at the predicted TG loci, including autoregulation of NR2F1 and TFAP2A themselves and the inferred regulation of both *NES* and *MEX3A* by both TFs. While NR2F1 and TFAP2A are also bound at the TSS and at putative intronic enhancers of *PRRX2* to regulate its expression (supported by high H3K27ac signals), NR2F1 appears to regulate the glutathione peroxidase *GPX7* only by binding to an enhancer 10 kb upstream of the promoter. Additional predicted TGs with corresponding NR2F1 and TFAP2A ChIP-seq tracks are shown in Supplementary Figure 8. Thus, our integrative time-resolved multi-omics analysis revealed transcriptional hierarchies, combinatorial TF regulation, and tightly controlled TGs driving neuroectodermal differentiation, NC specification, and human CNCC populations.

### Non-negative matrix factorization analysis infers core TFs in human CNCC differentiation and early lineage priming

To identify distinct regulatory signatures active at each timepoint, we applied NMF to the PECA2-inferred TRS matrices (Figure 2), which was previously shown to reveal subpopulation-specific regulation from bulk data [67]. The optimal number of signatures (k) per timepoint was determined by evaluating six performance metrics, resulting in factorization rank k=5 for d5, d8, d11 and k=7 for p2 (Supplementary Figure 9). The heatmap (Figure 3a) shows the transcriptional signatures identified at each timepoint, i.e. the contribution of a TF in a signature, of top ranked TFs per signature across timepoints. These contributions are similar to loadings in PCA, however, they are positive in NMF. We observed increasing importance of known CNCC regulators such as PRRX1/2, ALX1/4 and DLX1/2 [19] and decreasing relevance of anterior neuroepithelial TFs such as RAX, SOX2 or ZIC2/3 (Figure 3a) [57]. Signatures enriched for RAX and ZIC2/3 on d5-d11 but lower SOX2 exposure (d5_Sig2/3, d8_Sig3, d11_Sig2), likely represent a neural plate border like signal, the region, in which NC are induced [2]. In contrast, signatures with higher SOX2/3 exposure (d5_Sig4, d8_Sig2/4) may correspond to nascent NPC present in NECs [68]. NC specification between d5 and d11 (d8_Sig5, d11_Sig3), is characterised by high exposure of SOX10, confirmed on protein level (Supplementary Figure 1c-d). Subsequent downregulation of *SOX10* in p2 reflects a known feature of ectomesenchymal CNCC fate acquisition [69] (Supplementary Figure 1b, Figure 3a). Remarkably, p2 cells, which are generally more homogenous [22] still exhibit structured transcriptional heterogeneity. Namely, p2_Sig1/6 signatures display lower contribution of the ectomesenchymal CNCC regulators of the PRRX, ALX and DLX family and compared to e.g. HEY2. Since HEY2 is a neurogenic regulator also expressed in murine cranial ganglia [70,71], our finding positions this TF as a novel candidate regulator of neurogenesis from multipotent CNCCs in humans. TFs like SOX8 and PKNOX2 with high contribution in p2_Sig7 are implicated in ear morphogenesis [61,72]. Together with the tight regulation by key CNCC TFs (Figure 2d), the NMF findings strongly suggest PKNOX2 as an additional novel candidate for ectomesenchymal human CNCC differentiation. Besides these timepoint-specific TFs, we identified a substantial set of TFs with continuous contribution throughout differentiation, (e.g. CTCF, MYCN, MYBL2, GABPA, or TCF4/12) (Figure 3a). The contribution of these more general regulators of the transcription and proliferation [73–77] further confirm the dynamic processes underlying human CNCC differentiation. In addition to the differences across timepoints, we quantified the similarity of NMF-identified signatures across timepoints using Jaccard similarity analysis (Figure 3b) to address the recurring question of fate restriction timing in NC development. Our data are consistent with early fate priming. In particular, ectomesenchymal signatures (SOX4/8 and PKNOX2 in d8_Sig4 and p2_Sig7) and neuronal signatures (FOXD2, SIX2, PAX3 in d8_Sig1 and p2_Sig3) begin to be induced early in differentiation.

**Figure 3.**
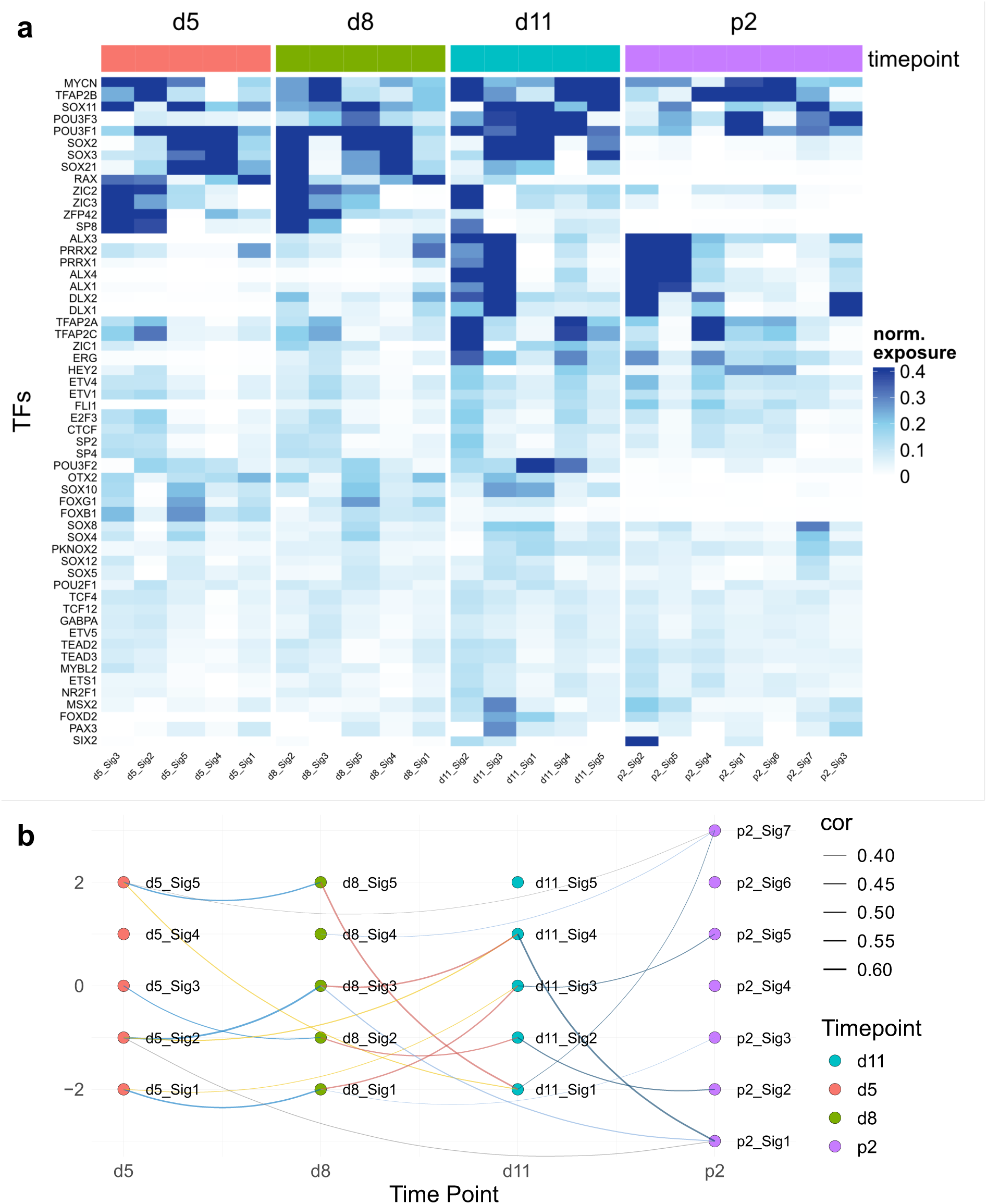
NMF analysis infers core TFs in human CNCC differentiation and early lineage priming. **a** TF contribution per signature and timepoint reveals timepoint-specific and continuously relevant TFs as shown for the optimal factorization ranks per time point (d5-11: k=5, p2: k=7). **b** Jaccard similarity between signatures per timepoint shows early lineage priming.

Altogether, our NMF analysis robustly captures the dynamic GRNs governing human CNCC differentiation, identifying conserved TFs and previously underappreciated TFs in the human system (HEY2 and PKNOX2). Moreover, we determined core regulatory signatures and and showed early establishment and progressive refinement of lineage-specific regulatory signatures as differentiation advances.

### Timepoint and cell type specific transcriptional regulation during human cranial neural crest development

Consistent with previous studies [67], our data suggest that the NMF-derived TF signatures reflect key cell populations and lineage biases essential for generating the diverse cell types required for proper cranial NC development. To validate this interpretation as distinct cell states, we performed single-cell RNA-sequencing (scRNA-seq) yielding 49,000 cells equally distributed across the timepoints. The Uniform manifold approximation and projection (UMAP) (Figure 4a) recapitulated the differentiation continuum of d5/d8/d11 observed in our bulk CAGE data (Figure 1c-d, Supplementary Figure 1b) with clearly separated p2 human CNCC. As shown in Figure 4b, early NECs at d5 homogeneously express NPC regulators such as *SOX2* with *PAX3* positive cells emerging, indicating a neural plate border-like population. Early migratory NC by d11 co-express *SOX9*, *TFAP2A*, and *TWIST1* and begin to acquire a cranial identity. Subsequent shared-nearest neighbour-based clustering revealed 11 distinct cell states (Figure 4c) with p2 diversifying into chondrogenic/osteogenic lineage marked by *MSX1* [78] and *CDH19* [79] in cluster 2 and the neuronal/glial linage marked by *NGF* [80] in cluster 4.

**Figure 4.**
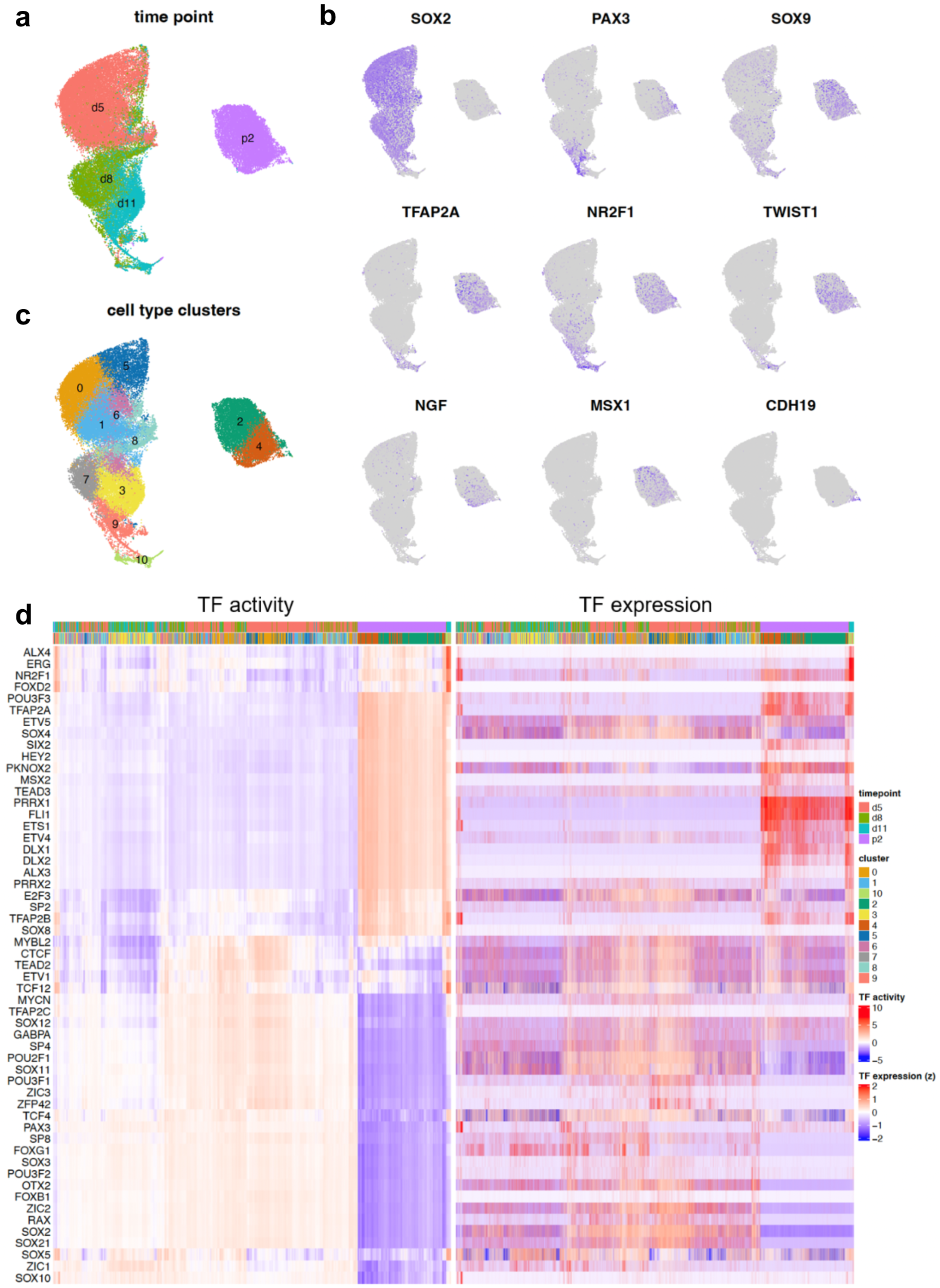
Identification of timepoint- and cell type-specific core regulators during human CNCC differentiation. **a** scRNA-seq dataset (n=4) encompassing d5, d8, d11, and p2 of the human CNCC differentiation. **b** Gene expression of selected markers confirms the differentiation continuum and lineage biases in the cranial NC which allows annotation of cell states in the clustered data (**c**). **d** Expression (right) and inferred activity (left) of core TFs obtained by NMF shows cell state-specific activity. Relative expression or activity across samples per gene is shown (z-scores).

To predict activity of the core TFs identified by NMF, we next integrated the PECA2-derived regulons from bulk data with the scRNA-seq profiles. We used top-ranking TGs per TF (based on TRS) and only retained those TF-TG pairs with significant correlations across the scRNA-seq dataset to construct TF regulons required for TF activity inference. TF activity clustered cells according to timepoint (Figure 4d, right heatmap), denoising sparse expression data [81,82]. SOX10 positive cells showed highest SOX10 activity, clustering together especially at d11 (see also Supplementary Figure 1d). ALX3/4, key craniofacial regulators, mutations of which cause frontonasal dysplasia (FND1/2) [83], displayed time-dependent activity: ALX4 peaks in early migratory CNCC, ALX3 in differentiated human CNCCs (Figure 4), possibly suggesting time point specific activity of these TFs underlying the distinct phenotypes of FND1/2. In agreement with the cell state specific expression and activity of NMF-identified TFs, their aggregated activity was also distinct among scRNA-seq subpopulations (Supplementary Figure 10). These data indicate that the *in vitro* human CNCC differentiation recapitulates several *in vivo* NC developmental intermediates [2,48] with timepoint- and subpopulation-specific core TF expression/activity. Thus, our unique time-resolved single-cell map of human CNCC development serves as a powerful tool to explore cell state-specific processes in both health and disease.

### Network-based linkage of OFC-associated variants to known and novel risk genes

Having confirmed that the *in vitro* protocol recapitulates several aspects of tissue diversity and GRNs relevant to craniofacial morphogenesis in a human system, we queried whether human genetic variation associated with the common craniofacial condition nsCL/P might act through these GRNs. In particular, we leveraged the GRNs to link nsCL/P-associated variants to the TGs they control to prioritize nsCL/P risk genes at associated loci. Therefore, we used 45 lead risk loci from a recent nsCL/P meta-analysis [28] within windows of high linkage disequilibrium (LD; r^2^ > 0.6). Of the 45 loci, 29 partially overlapped with our PECA2 REs across all timepoint-specific GRNs (significant enrichment based on permutation test, p = 0.002; Supplementary Figure 11), identifying them as candidates that may disrupt REs and modulate TG expression particularly in CNCCs. Based on the GRNs, these 29 loci were linked to 70 putative TGs (Table 1). The predicted TGs include well-known links between nsCL/P risk regions and genes implicated in CNCC development and orofacial clefting, e.g. *PAX7*, *COL8A1*, *FILIP1L*, *GADD45G* [84], *FGFR1* [85,86], and *PTCH1* associated with Hedgehog signaling [87], or *KRT8* [88].

**Table 1.** nsCL/P-associated SNP-TG links according to the PECA2 predictions. Lead variants and their window of high LD in a European population are shown with the predicted TG controlled by REs falling within the high-LD windows per variant. Predicted TGs outside of the LD window (r^2^ > 0.6) are indicated with an asterisk.

| SNP | chr | LD $r^2 > 0.6$ start | LD $r^2 > 0.6$ end | TG |
| --- | --- | --- | --- | --- |
| rs742071 | chr1 | 18638566 | 18666089 | *PAX7* |
| rs34746930 | chr1 | 19369502 | 19458400 | *CAPZB* |
| rs560426 | chr1 | 94075118 | 94092554 | *FNBP1L* * |
| rs560426 | chr1 | 94075118 | 94092554 | *ABCA4* |
| rs642961 | chr1 | 209667927 | 210300043 | *SYT14* |
| rs642961 | chr1 | 209667927 | 210300043 | *CAMK1G* * |
| rs642961 | chr1 | 209667927 | 210300043 | *C1orf74* |
| rs287982 | chr2 | 9819981 | 9901155 | *GRHL1* * |
| rs287982 | chr2 | 9819981 | 9901155 | *MBOAT2* * |
| rs6740960 | chr2 | 41954538 | 41971077 | *PKDCC* * |
| rs7590268 | chr2 | 43274445 | 43609877 | *LRPPRC* * |
| rs7590268 | chr2 | 43274445 | 43609877 | *DYNC2LI1* * |
| rs7590268 | chr2 | 43274445 | 43609877 | *PLEKHH2* * |
| rs7590268 | chr2 | 43274445 | 43609877 | *SLC3A1* * |
| rs7590268 | chr2 | 43274445 | 43609877 | *HAAO* * |
| rs793464 | chr3 | 99704428 | 100195985 | *TFG* * |
| rs793464 | chr3 | 99704428 | 100195985 | *CMSS1* |
| rs793464 | chr3 | 99704428 | 100195985 | *LNP1* * |
| rs793464 | chr3 | 99704428 | 100195985 | *COL8A1* |
| rs793464 | chr3 | 99704428 | 100195985 | *FILIP1L* |
| rs1907989 | chr4 | 4772239 | 4819003 | *STK32B* * |
| rs1907989 | chr4 | 4772239 | 4819003 | *CYTL1* * |
| rs908822 | chr4 | 123866230 | 124053159 | *ANKRD50* * |
| rs6449957 | chr5 | 68147564 | 68224398 | *PIK3R1* |
| rs13317 | chr8 | 38162889 | 38453392 | *FGFR1* |
| rs13317 | chr8 | 38162889 | 38453392 | *EIF4EBP1* * |
| rs13317 | chr8 | 38162889 | 38453392 | *LETM2* |
| rs13317 | chr8 | 38162889 | 38453392 | *ASH2L* * |
| rs12543318 | chr8 | 87856111 | 88012602 | *MMP16* * |
| rs957448 | chr8 | 94347409 | 94615738 | *ESRP1* * |
| rs957448 | chr8 | 94347409 | 94615738 | *RAD54B* |
| rs10908902 | chr9 | 89565600 | 89629605 | *S1PR3* * |
| rs10908902 | chr9 | 89565600 | 89629605 | *GADD45G* |
| rs10512248 | chr9 | 95454593 | 95516362 | *PTCH1* |
| rs3758249 | chr9 | 97810881 | 97913694 | *GALNT12* * |
| rs3758249 | chr9 | 97810881 | 97913694 | *TRIM14* * |
| rs3758249 | chr9 | 97810881 | 97913694 | *CORO2A* * |
| rs7078160 | chr10 | 116881884 | 117137256 | *ENO4* |
| rs7078160 | chr10 | 116881884 | 117137256 | *VAX1* |
| rs3741442 | chr12 | 52946815 | 52963551 | *KRT8* |
| rs705704 | chr12 | 55975721 | 56088396 | *GDF11* * |
| rs705704 | chr12 | 55975721 | 56088396 | *TAC3* * |
| rs705704 | chr12 | 55975721 | 56088396 | *IKZF4* |
| rs705704 | chr12 | 55975721 | 56088396 | *ERBB3* |
| rs705704 | chr12 | 55975721 | 56088396 | *MYL6* * |
| rs705704 | chr12 | 55975721 | 56088396 | *RAB5B* |
| rs705704 | chr12 | 55975721 | 56088396 | **SUOX** |
| rs705704 | chr12 | 55975721 | 56088396 | **CNPY2** |
| rs705704 | chr12 | 55975721 | 56088396 | **CDK2** |
| rs705704 | chr12 | 55975721 | 56088396 | **RPS26** |
| rs705704 | chr12 | 55975721 | 56088396 | **MYL6B** |
| rs2304269 | chr12 | 71563011 | 71721887 | **THAP2** |
| rs2304269 | chr12 | 71563011 | 71721887 | **RAB21\*** |
| rs1873147 | chr15 | 63019226 | 63033349 | **TPM1\*** |
| rs1873147 | chr15 | 63019226 | 63033349 | **TLN2\*** |
| rs28689146 | chr15 | 74419836 | 74737066 | **CSK\*** |
| rs28689146 | chr15 | 74419836 | 74737066 | **ARID3B** |
| rs28689146 | chr15 | 74419836 | 74737066 | **EDC3** |
| rs28689146 | chr15 | 74419836 | 74737066 | **IMP3\*** |
| rs28689146 | chr15 | 74419836 | 74737066 | **CYP1A1** |
| rs8049367 | chr16 | 3913185 | 3945230 | **HCFC1R1\*** |
| rs9891446 | chr17 | 9028764 | 9044391 | **NTN1** |
| rs1838105 | chr17 | 46899673 | 47096919 | **GOSR2** |
| rs1838105 | chr17 | 46899673 | 47096919 | **SP2\*** |
| rs1588366 | chr17 | 62966043 | 63343461 | **DCAF7\*** |
| rs3746101 | chr19 | 2038819 | 2049315 | **MKNK2** |
| rs3746101 | chr19 | 2038819 | 2049315 | **OAZ1\*** |
| rs3746101 | chr19 | 2038819 | 2049315 | **MOB3A\*** |
| rs3091552 | chr20 | 46811366 | 46927877 | **SLC12A5\*** |
| rs3091552 | chr20 | 46811366 | 46927877 | **EYA2** |

Besides confirming known risk genes located within associated regions, the time-resolved GRNs uncovered distal variant-gene associations, mapping 40 out of the 70 risk genes that lie outside of the LD region of the respective linked variant (Table 1, asterisk). Our approach not only successfully identified the distal regulation of *PKDCC* by rs6740960, an interaction previously validated in chondrocytes, [89], but also uncovered several novel distal variant-to-gene relationships. These are of particular relevance, as they reveal novel links between well-established orofacial clefting genes, including *MMP16* [90], *ESRP1* [91] or *TPM1* [92] and known distal regulatory elements (REs) for the first time. The value of these novel interpretations can be exemplified by the well-studied nsCL/P-associated *IRF6* locus. Here, the lead SNP rs642961 resides in a putative enhancer region [93], although the causal variant within the high-LD window remains to be determined. Our data indicate that the high-LD window around rs642961 overlaps a RE predicted to control distal genes *SYT14*, *C1orf74* and *CAMK1G*, but not the nearby genes *IRF6* or *UTP25* (Supplementary Figure 12a). Consistent with our findings, a Chinese cohort study identified nsCL/P-associated variants at the SYT14 locus that show only modest linkage disequilibrium (r² < 0.4) with IRF6 SNPs [94]. Although SYT14 has not been extensively studied in CNCC differentiation, our data suggest it may represent a relevant candidate gene contributing to neural crest involvement in nsCL/P etiology.

Another candidate is rs7590268, located at the *THADA* locus, [95]. While *THADA* is expressed throughout human CNCC differentiation (Supplementary Figure 12d), functional experiments have so far not revealed a role of the gene in orofacial development [96]. Instead, our data indicate that REs in LD with rs7590268 may regulate *LRPPRC*, *DYNC2LI1*, *SLC3A1*, and *HAAO* (Supplementary Figure 12c), genes implicated in craniofacial development. *LRPPRC* mutations are linked to French-Canadian Leigh disease characterized by craniofacial dysmorphism without orofacial clefting, besides several other clinical features [97]. In contrast, *DYNC2LI1* mutations are associated with a complex clinical phenotype including orofacial clefts [98]. Albeit sparsely expressed throughout human CNCC differentiation (Supplementary Figure 12c), *DYNC2LI1* is responsible for maintaining the primary cilia structure and proper Hedgehog signaling in many cell types, including neural crest cells, making it a prime candidate TG for the RE containing the rs7590268 variant. *PLEKHH2*, which shows low and heterogeneous expression during the differentiation (Supplementary Figure 12c), is predicted to be actin binding but only related proteins like PLEKHA5 and PLEKHA7 have been implicated in nsCL/P in distinct cohorts [99,100]. *SLC3A1* is only lowly expressed at all time points of differentiation (Supplementary Figure 12d), however, homozygous deletions near or encompassing *SLC3A1* are associated with distinct craniofacial features [101]. The fifth predicted TG for a RE at the OFC risk locus around rs7590268 is *HAAO* (Supplementary Figure 12c-d), which shows sparse expression in p2. While its role in craniofacial morphogenesis remains poorly understood, a homozygous knockout mouse exhibited a cleft palate [102], suggesting *HAAO* as a potential OFC susceptibility gene. Thus, REs containing rs7590268 may control multiple TGs important in orofacial morphogenesis, suggesting combined dysregulation of several of these predicted TGs at this locus in nsCL/P.

Overall, using the time-resolved GRN, we were able to map nsCL/P SNPs to known risk and novel candidate genes that expand the distal regulatory relationships between known variants and risk genes for nsCL/P. In particular, we revealed both context-specific regulatory effects, such as the *IRF6* locus likely acting through *SYT14* in CNCCs, as well as loci with potentially complex multi-gene regulation, as illustrated by the *THADA* locus.

To assess whether the expression of the 70 putative TGs is particularly enriched in sub-populations of the CNCC differentiation, thereby identifying the cellular context most relevant for nsCL/P pathogenesis, we performed polygenic enrichment analysis and single-cell disease relevance score (scDRS) analysis [103]. Leveraging our complementary single-cell RNA-seq data generated at the same four timepoints of the CNCC differentiation, we identified significant cluster-trait associations in cluster 2 in p2 (Figure 4b) with an ectomesenchymal (*MSX1*/*CDH19*-positive) bias (Figure 5a-b). This finding suggests that within the NC compartment, nsCL/P risk gene expression may be particularly enriched in CNCCs already biased towards an osteo/chondrogenic fate, complementing previously described disease-relevant mechanisms in other cell types such as the periderm [104].

**Figure 5.**
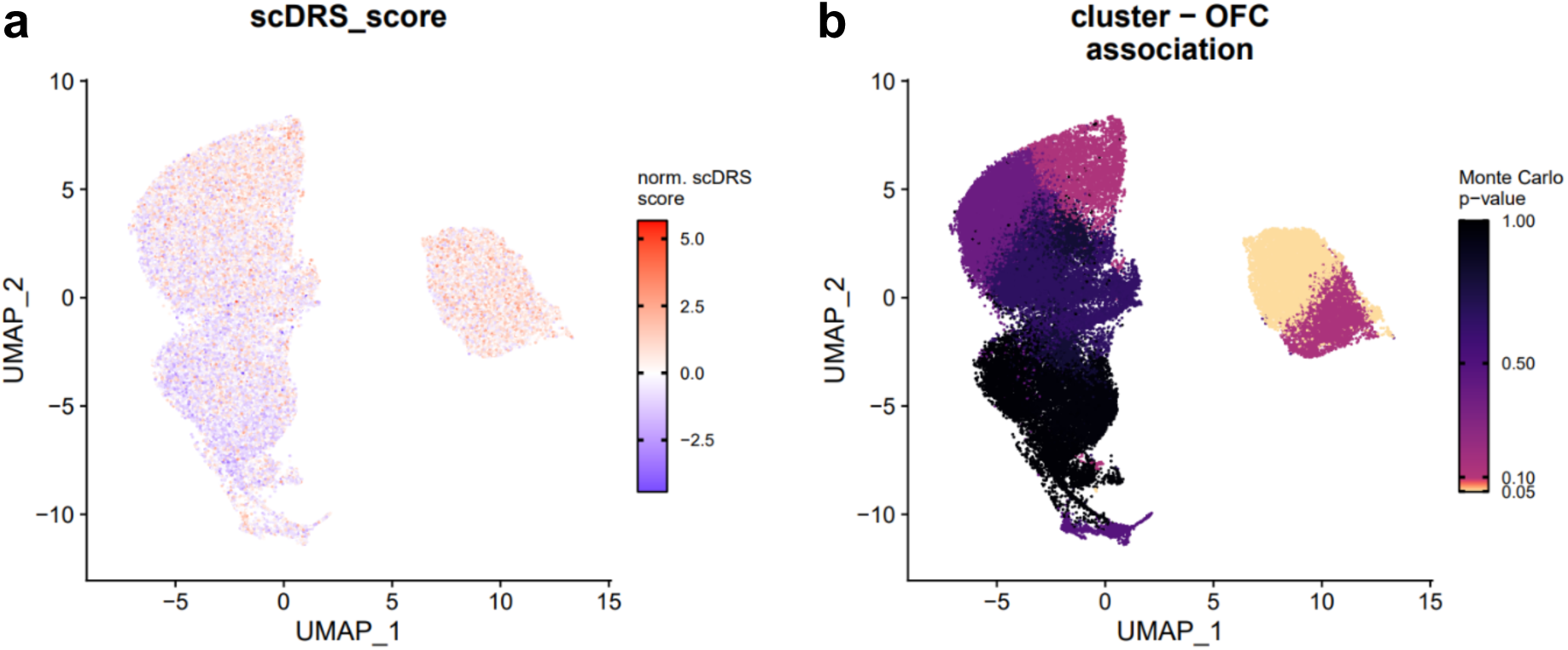
Single-cell disease relevance score analysis reveals association of ectomesenchymally biased human CNCC with nsCL/p. **a** scDRS calculated from the 70 PECA2-inferred nsCL/P risk genes. **b** Significant cluster-trait association for the chondrogenesis and osteogenesis-biased human CNCC.

Taken together, the time-resolved GRN analysis enabled the linking of nsCL/P-associated variants to both known and novel target genes and predicted distal (beyond LD) as well as context-specific (ectomesenchyme-biased) regulatory effects, expanding the interpretation of non-coding variants. In addition, data generated in this study may help prioritize further genetic variants in nsCL/P and neurocristopathies more broadly.

## Discussion

Although interpreting non-coding variants within dynamic craniofacial GRNs is crucial for understanding common malformations like the highly prevalent neurocristopathy nsCL/P, a human-specific time-resolved multi-omics framework for CNCC development is lacking. Previous work on NC gene regulation is mainly based on model organisms [2,4,5] or on snapshots of later stages of this process [19]. Hence, to our knowledge, our transcriptomic and chromatin accessibility profiling provides the first comprehensive GRNs with high temporal resolution that dynamically govern early human CNCC differentiation, including the neuroepithelial fate, NC induction, specification and the acquisition of a cranial identity.

Mapping nsCL/P-associated non-coding variants onto the GRNs identified 29 out of 45 variants [28] and linked them to 70 putative TGs, including both known variant-target gene links for e.g., *FGFR1*, *COL8A1* and *PTCH1* and previously not reported links for e.g., *SYT14* and *TPM1*. We speculate that the remaining 16 out of 45 nsCL/P risk loci [28] may affect non-CNCC cell types contributing to craniofacial morphogenesis, such as the periderm [107].

Although the predicted TGs have been previously linked to craniofacial development, they have not been associated with these nsCL/P variants. Moreover, 40 of the predicted 70 TGs lie outside the risk loci (LD), indicating novel putative distal regulatory relationships rather than a simple link to the nearest gene. These findings broaden our knowledge of the genetic basis of nsCL/P and demonstrate the power of time-resolved GRNs for discovering relationships in human craniofacial biology that would be missed in analyses based on a single timepoint. For example, despite the identification of variants associated with nsCL/P at the *THADA* locus [95], the gene itself has not been implicated in orofacial morphogenesis [96]. Our data suggest that this nsCL/P risk locus may instead affect other genes surrounding this region, e.g. *HAAO*. In addition, our data indicate that variants at the *IRF6* locus modulate the expression of genes other than *IRF6* in human CNCC, particularly *SYT14*.

Of note, the expression of the of the TGs newly linked to distal nsCL/P risk variants is generally enriched in the chondrogenic/osteogenic human CNCC subpopulation, which is significantly associated with the condition. These findings therefore contribute to a better subtype resolution of previously described cell type nsCL/P-associations, which implicated the cranial epithelium and a pharyngeal arch population [108]. More exactly, the data indicate that the pharyngeal arch CNCCs with an already acquired ectomesenchymal bias are responsible for this association, in line with their future contribution to orofacial morphogenesis [10]. Complementing the bulk CAGE and ATAC-seq data with scRNA-seq profiles revealed ectomesenchymally biased human CNCCs as a susceptible subpopulation in nsCL/P pathogenesis, thereby narrowing down the cellular context e.g., for future functional validation of non-coding genetic variants. Taken together, our work demonstrates that the time-resolved network-based interpretation of genetic variants associated with congenital conditions or birth defects substantially improves the prediction of the relevance of the mapped non-coding nsCL/P variants by refining the variant-to-gene relationships. Ultimately, the data provide a valuable resource for future, e.g., CRISPR-based screening to functionally test the transcriptional hierarchies and the contribution of the newly suggested variant-to-gene relationships to congenital conditions like nsCL/P.

Overall, the integrative time-resolved multi-omic data demonstrate that several features of *in vivo* NC development can be reproduced in the hiPSC-based differentiation protocol [1,48], including a continuum of NC induction and specification from the neuroepithelium and chondrogenic/osteogenic and neuronal fate biases. Thus, our data position *in vitro* differentiation as a valuable orthogonal approach to human embryonic developmental studies, which are difficult to access. Our framework enabled us to reveal core, timepoint-related core TFs for each of these differentiation stages and confirmed the network inferences for NR2F1 and TFAP2A by corresponding ChIP-seq data, highlighting the accuracy of our inferences. Unbiased identification of core TFs by NMF of the GRNs revealed many conserved regulators, including neuroepithelial regulators of the SOX and ZIC families, SOX10 at NC specification of migratory and cranial regulators of the PRRX, ALX and DLX families of TFs [2,4,5]. In addition, some TFs that have previously not been studied in detail in NC biology, particularly in the human system, were among the inferred core TFs. Although the FOX family of TFs, for example, has been strongly implicated in different stages of NC development [105], FOXD2 in particular has remained largely uncharacterised. So far, its expression had only been observed in CNCCs around the eye and in the ethmoidal and mandibular processes [106]. Our data, however, suggest that FOXD2 acts as an important regulator in early migratory human CNCCs, particularly at day 11 and in p2.

We are cautious in interpreting absolute regulatory hierarchies as suggested by the networks. Here, *ALX3*, for example, was not strongly regulated by either NR2F1 or TFAP2A in p2, although both were bound to an enhancer at the *ALX3* locus in d11 human CNCC [20]. At this point we cannot ascertain whether the transcriptional regulatory input received by *ALX3* may be different between d11 and p2 human CNCC, highlighting the need for orthogonal data modalities, such as additional ChIP-seq data, for each time point to corroborate the inferences. In addition, increasing temporal resolution may further refine these regulatory relationships. Likewise, the continuous improvement of single-cell multi-omic approaches [109,110] will help resolve the regulatory relationship underlying these cell state changes better in the future.

Together, the time-resolved multi-omic analysis of human CNCC differentiation illuminates the core regulatory relationship of healthy human CNCC development and establishes network-based variant interpretation as a useful framework for linking non-coding genetic variation to cell-type-specific developmental pathology.

## Methods

### hiPSC cell culture and cranial neural crest cell differentiation

hiPSC cells were cultured in StemMACS iPS-Brew XF (130-104-368, Miltenyi Biotec) on Geltrex (A1413202, Thermo Fisher) coated tissue culture dishes kept at 37 °C, 5% CO_2_. The cells were routinely passaged using Versene (15040066, Thermo Fisher) once they reached a confluency of 70%-80%. For the differentiation of the hiPSC in human CNCCs following published protocols [23], the hiPSC cells were grown until they reached full confluency The colonies were detached from the tissue culture dishes by adding fresh Collagenase IV (17104019, Thermo Fisher) diluted in KnockOut-DMEM (10829018, Thermo Fisher) to a final concentration 3 µg/ml per well. The plate was incubated at 37 °C for 1-2 hours until the colonies detached from the plate. Upon detachment of the colonies, medium sized aggregates of the colonies were generated by manually disturbing the colonies using a P1000 and transferring the cell aggregates into a petri dish containing human CNCC differentiation medium (1:1 ratio Neurobasal (21103-049, Gibco) media with DMEM-F12 + Glutamax (31331-028, Gibco) supplemented with 0.5x B27 (17504-044, Gibco), 0.5x N2 (17502-048, Gibco), 0.5x Glutamax 100x (35050038, Thermo Fisher), 20 ng/ml bFGF (100-18B, Thermo Fisher), 5 µg/ml insulin (11376497001, Sigma-Aldrich), 20 ng/ml EGF (AF-100-15-500UG, Thermo Fisher). During the 11 day differentiation protocol, neuroectodermal spheres (NECs) form and attach to the petri dish to give rise to human CNCCs. Medium changes were performed routinely. Upon initial attachment of the NECs usually around day 5 of the differentiation protocol. At day 11 adherent CNCCs were harvested by detaching them from the petri-dish with Accutase (A6964, Sigma-Aldrich) and transferring them into tissue culture dishes coated with 5 µg/ml fibronectin (FC010, Merck) containing NC maintenance medium (1:1 ratio Neurobasal Media with DMEM-F12 + Glutamax supplemented with 0.5x B27, 0.5x N2, 0.5x Glutamax 100x, 20 ng/ml bFGF, 1 mg/ml BSA (11020-021, Thermo Fisher), 20 ng/ml EGF).

### Cap analysis of gene expression

CAGE samples were acquired from four time points of the *in vitro* human CNCC differentiation protocol starting from hiPSCs [22,44], namely d5, d8, d11, and d16/17, corresponding to the confluent cell population of human CNCCs after the second passage of the d11 NCCs (p2). Three replicates were generated for each time point and total RNA extracted with PureLink RNA Mini Kit (12183018A, Thermo Fisher) including on-column DNA digest with the PureLink DNase Set (12185010, Thermo Fisher). All samples were quality checked by automated gel electrophoresis of 2,5 ng per replicate with the RNA 6000 Pico kits (5067-1513, Agilent) on a 2100 Bioanalyzer.

CAGE libraries were constructed as outlined in the SLIC-CAGE protocol by [111]. Specifically, 2500 ng of total RNA from each replicate were mixed with 2500 ng of prepared SLIC-CAGE carrier RNA. Using these carrier-combined samples as starting material, first-strand cDNA synthesis was performed. The produced RNA:cDNA hybrids were subjected to oxidation to expose the terminal RNA nucleotides, notably the 7-methylguanosine (m7G), which permitted biotinylation of only capped RNA species. Overhanging RNA segments that are not part of the RNA:cDNA hybrids were degraded using RNase ONE digestion to guarantee that the subsequent pull-down with paramagnetic Streptavidin beads (i.e., cap-trapping) exclusively isolates the biotinylated m7Gs. cDNAs derived solely from capped RNAs were relesed from beads and hybrids via combined RNaseH and RNase ONE digestion, the Illumina-compatible 5’ and 3’ nAnT-iCAGE linkers were ligated, and second-strand cDNA was synthesized. Carriers were subjected to digestion, the generated library was size selected and underwent PCR amplification. The average library length was assessed based on the fragment distribution determined on the 2100 Bioanalyzer with the High Sensitivity DNA kit (2067-4626, Agilent) and quantified with the Quant-iT PicoGreen dsDNA kit (P7589, Thermo Fisher). Equimolar pooling of sets containing 8 samples was carried out before single-end sequencing on a NextSeq550 Illumina sequencer with High Output v2.5 (75 cycles) reagents (20024906, Illumina) employing standard Illumina primers. The NextSeq PhiX Control Kit from Illumina was used to spike the libraries with 3% PhiX (FC-110-3002, Illumina).

Libraries were demultiplexed by SLIC-CAGE adapter barcodes, reads trimmed to remove linker sequences and FASTX-Toolkit used to filter for a minimum sequence quality of Q30 in at least 50% of read sequence. rRNAdust was employed to remove reads matching subsequences of the human ribosomal DNA complete repeating unit (U13369.1, [112]). BWA was used to map the data to the hg38 assembly of the human reference genome while permitting a maximum edit distance of 2. Transcription start sites were identified from the mapped reads and the signal from plus and minus strand separated. Downstream CAGE data analysis was performed according to the CAGEfightR workflow [35,113]: Briefly, CAGE-TSSs (CTSSs) mapping to overlapping regions were quantified and then clustered. Unidirectional CTSS clusters correspond to putative TSSs and bidirectional CTSS clusters correspond to putative enhancers. TSSs with at least one count in at least four samples were retained. Enhancers with at least one count in at least three samples were retained. We further discarded enhancers that did not map to annotated intergenic or intronic regions [114]. The TSS signal overlapping annotated genes was summed to obtain gene expression levels [35]. The gene expression data and the eRNA data was normalized to library size separately, subjected to principal component analysis (PCA) as well as differential expression analysis using DESeq2 [115].

### ATAC-seq data

ATAC-seq was performed on the corresponding samples as for the generation of the CAGE data described above. Cell samples were lysed by osmotic pressure to release intact nuclei. For this, the cells were centrifuged at 1000 rpm for 2 min and the pellet resuspended in 500 µl hypotonic lysis buffer (10mM Tris-HCl, ph7.4; 10mM NaCl; 3mM MgCl2; 1% v/v BSA; 0,1% v/v NP-40 Substitute (11754599001, Roche); 0,1% v/v Tween20 (11332465001, Roche); 0,01% v/v Digitonin (G9441, Promega); 4 mU/µl SUPERase·In (AM2696, Thermo Fisher); 4 mU/µl Protector RNase Inhibitor (3335402001, Sigma-Aldrich)). After lysing cells for 5 min on ice, the entire solution was diluted with 9,5 ml hypotonic resuspension buffer (10mM Tris-HCl, ph7.4; 10mM NaCl; 3mM MgCl2; 1% v/v BSA; 0,1% v/v Tween20 (655206, Sigma-Aldrich); 4 mU/µl SUPERase·In (AM2696, Thermo Fisher); 4 mU/µl Protector RNase Inhibitor (3335402001, Sigma-Aldrich)). Nuclei were separated from cell debris by centrifuging the samples in 15 ml Falcon tubes for 10 min at 4°C and 500xg. After discarding the debris-containing supernatant, nuclei were resuspended in 300 µl hypotonic resuspension buffer and counted with a Bürker cell chamber. For each replicate the volume corresponding to 100k nuclei was separated and centrifuged at 4°C and 500xg for 10 min. After discarding the supernatant, the individual nuclei pellets were resuspended in each 20 µl of tagmentation reaction (10 µl 2x Tagment DNA buffer (15027866, Illumina); 2 µl ddH_2_O; 6.6 µl 1x PBS; 0,2 µl 10% Tween20 (655206, Sigma-Aldrich); 0,2 µl 1% Digitonin (G9441, Promega); 1 µl TDE1 enzyme (15027916, Illumina). These tagmentation reactions were incubated for 30 min at 37°C and 550 rpm on a thermomixer. DNA was extracted with the MinElute Reaction Cleanup kit (28204, Qiagen) and eluted in 10 µl Buffer EB (19086, Qiagen). The entire eluted, transposed DNA was mixed with 10 µl ddH_2_O, 25 µl NEBNext High Fidelity 2x PCR Master Mix (M0541, NEB) and each 2,5 µl of the custom Nextera i5 and i7 adapters as described in [116]. These were amplified using the following PCR conditions: 72°C for 5 min, 98°C for 30 sec, followed by 10 cycles of 98°C for 10 sec, 63°C for 20 sec and 72°C for 1 min. The resulting library was cleaned by double-sided AMPure XP bead (A63880, Beckman Coulter) purification that removes primer dimers and fragments surpassing a length of 1kbp. The quality and average fragment lengths of the resulting libraries was assessed on the 2100 Bioanalyzer with the High Sensitivity DNA kit (2067-4626, Agilent) and the libraries quantified with the Quant-iT PicoGreen dsDNA kit (P7589, Thermo Fisher). Multiplexed libraries were sequenced paired-end on a NextSeq550 Illumina sequencer with High Output v2.5 (150 cycles) reagents (20024907, Illumina) employing standard Illumina primers. The multiplexed library was spiked with 3% PhiX (FC-110-3002, Illumina).

The sequenced raw files were pre-processing including quality control, alignment, deduplication, peak calling and quantification of the ATAC-seq signal using the standard ENCODE pipeline [117]. The differential accessibility analysis workflow used here makes use of the csaw package [118], similar to [119]. Briefly, we extracted the optimal peak set per sample according to overlap reproducibility (.narrowPeak files) and created the union of peaks across all samples to recount the deduplicated reads (.bam files) for these windows. Parameters for re-counting were set to max.frag=1000, pe=“none” This yielded a count matrix for the ATAC-seq data. We only retained genomic regions where at least one sample had more than 10 counts. The count matrix was normalized to library size and subjected to PCA and differential accessibility analysis with DESeq2 [115].

### Time-course analysis of chromatin accessibility, gene and eRNA expression

Genes and eRNAs with variable time-course expression as well as genomic regions with variable accessibility, i.e. a minimum log2-fold change of 1 when comparing each time point to any other, were included for the clustering of genes and eRNAs with similar expression dynamics and genomic regions with similar accessibility dynamics. This was performed using fuzzy c-means clustering using the Mfuzz package [120] with m=1.5 and the number of clusters (c) chosen per data modality as the highest number of clusters that did not produce two clusters of highly similar overall trajectories. The gene / eRNA expression data and chromatin accessibility data was first averaged across the three replicates for this analysis and then z-scored per gene or genomic region. Cluster-wise gene ontology (GO) enrichment analysis for the gene expression clusters was performed using the compareCluster() function from clusterProfiler [121] and querying all ontologies (biological process, molecular function, cellular component). P-values were adjusted with the Benjamini-Hochberg correction. Cluster-wise Genomic Regions Enrichment of Annotations Tool (GREAT) analysis was performed for the putative enhancer regions and differentially accessible regions using the rGREAT package [122]. GREAT analysis was limited to genomic regions that represented the overall trajectory of the time-course clusters best, disregarding regions with the 25% lowest cluster membership per clsuter. For GREAT analysis, we used only the human biological process (BP) ontology resource. Putative enhancer regions and differentially accessible regions per time-course cluster were subjected to known TF motif enrichment analysis with HOMER [51]. We used the findMotifsGenome.pl script with default options (-size 200 -mask) and specified the human genome build (hg38, v6.4). Results were loaded into R using the marge package [123] and top enriched known motifs per cluster sorted according to p-value, which was then visualized using the geom_tile() function from ggplot2. Gene and eRNA expression boxplots were obtained with the ggboxplot() function of the ggpubr package [124] built on top of ggplot2 [125]. Genome browser-like visualizations of the ATAC-seq data in the time-course were prepared with the Gviz package [126] and transcript annotation from TxDb.Hsapiens.UCSC.hg38.knownGene [114].

### Gene regulatory network inference and network analysis

GRN inference per time point using paired gene expression and chromatin accessibility data from each time point was performed using PECA2 [19,127]. According to the input requirements for PECA2 we normalized the counts from gene-level quantification from the CAGE-data to transcripts per million (TPM) and averaged the TPM values across the three replicates per time point. The aligned reads from the ATAC-seq data for the three replicates per time point were combined by concatenating the .bam-files in line with the recommendations by [19,127]. PECA2 was executed separately for each time point with the function call PECA.sh hg38. Output trans-regulation score (TRS) matrices with TFs as rows and TGs as columns (TFTG_score_norm.txt) were loaded into R for further analysis and visualization. A .csv file was prepared containing the TRS for three TGs for the core TFs with highest TRS for subnetwork visualization in Cytoscape [128]. For this, the TRS for the five TFs per time point as possible TGs was included as well. Edges with TRS lower than the lower quartile of these TRS were discarded under the assumption that they would not correspond to true regulatory interactions. Time-course regulatory analysis including non-negative matrix factorization (NMF) of the TRS matrices per time point to obtain core regulatory modules was performed with ButchR [129]. The optimal number of signatures per timepoint was determined using optimal values for metrics such as cophenetic distance or silhouette score. Each signature is defined by the contribution or loading of the transcription factors to this signature. In NMF, these contributions are positive or zero values. NR2F1, TFAP2A and H3K27ac ChIP-seq tracks from d11 CNCC for predicted TGs of NR2F1 and TFAP2A originate from [20] and were visualized in the UCSC Genome Browser [130].

### scRNA-seq and data analysis

For d5, d8, d11 and p2 samples of the human CNCC differentiation (see hiPSC cell culture and cranial neural crest cell differentiation), scRNA-seq data was generated. Starting the dissociation, NECs/CNCCs were rinsed 2x with PBS, followed by Accutase incubation at 37°C. After 5 minutes, single-cell dissociation was checked, using a P1000 pipette. If clumps were observed after this 5-minute incubation period, the Accutase incubation was prolonged until the NECs were in a single-cell suspension without any visible clumps present. To inhibit further dissociation of the samples upon single-cell generation, the reaction was stopped using the respective growth medium of the cells. Single-cells were then pelleted at 300xg for 5 minutes at 4°C and resuspended in 250 µL PBS + 0.04% BSA. Following this, cells were first filtered using a 70 µm cell strainer, followed by an additional filtration step through a 40 µm cell strainer. Finally, cell viability was determined using the Cell Drop AO/PI Viability Assay, (DeNovix, # 31CD-AO-PI-1.5) according to the manufacturer’s instructions. If the viability was below 90 % an additional enrichment step was performed using the Dead Cell Removal Kit (Miltenyi Biotech, #130-090-101) according to the manufacturer’s recommendations.

Following the dissociation, cells were immediately processed using the Evercode^TM^ Cell Fixation v2/v3 (Pares Biosciences, #ECF3300) protocol for single-cell RNA sequencing according to the manufacturer’s instructions and cell numbers determined using a DeNovix CellDrop counting device. Paired-end sequencing was conducted at the Deep Sequencing Core Facility in Heidelberg with the sequencing parameters as recommended by Parse in the protocol.

The .fastq-files were converted to a count matrix using the official company pipeline according to the company instructions (Parse Biosciences). All R1 and R2 reads from different lanes were concatenated in one .fastq file, respectively. The indexed hg38 reference genome was generated according to company instructions and the count matrix obtained through the wrapper split-pipe function with settings –mode all –chemistry v2 / v3. The two count matrices were loaded into R, and processed further using standard Seurat workflows [131,132]. Time points were assigned according to the barcoding well 1. Datasets originating from the two different sequencing chemistries were integrated using SCTransform(). Clustering was performed with the FindNeighbors() function using the first 10 principal components followed by the FindClusters() function with a resolution of 0.4. TF activity inference was performed using the decoupleR package [82] with the run_ulm() function and minsize = 5. As regulon resource we derived tissue-specific regulons from the PECA2 GRNs. For this, the Pearson Correlation Coefficient (PCC) for expression of all TF-TG pairs with a TRS > 20 at any time point was determined, and only those with |PCC| > 0.1 were retained. Positive values were assigned a mode of regulation (mor) 1, negative values a mor of -1, corresponding to possible activation or repression, respectively.

### Mapping nsCL/P-associated variants to putative target genes

All putatively identified REs across the four GRNs were extracted and overlap assessed with genetic variants associated with nsCL/P. Specifically, we used the 45 lead SNPs published in a meta-analysis for nsCL/P-associated variants [28]. We took the window of high linkage disequilibrium (LD r^2^ > 0.6) per lead SNP as provided in the original publication, performed a lift-over from hg19 to hg38 using the liftOver() function from rtracklayer [133] and used these regions for overlap analysis with REs in the GRNs. Overlaps were identified with the findOverlaps() function from the GenomicRanges package [134]. To assess whether the overlap between nsCL/P-associated LD regions and GRN-derived REs exceeded what would be expected by chance, we performed a permutation test using the overlapPermTest() function from the regioneR package [135], randomizing the LD regions 1000 times across the hg38 genome with the alternative hypothesis that the observed overlap is greater than expected.

### Single-cell disease relevance score analysis

To assess the enrichment of nsCL/P genetic risk in our time-course scRNA-seq data of the human CNCC differentiation, we applied scDRS [136] to the integrated Seurat object exported as an .h5ad file. As the gene set input, we used the putative target genes of nsCL/P-associated regulatory elements identified through GRN overlap analysis (see Mapping nsCL/P-associated variants to putative target genes), formatted as a .gs file. scDRS disease relevance scores were computed with the compute-score function using raw counts, with data filtered to matching genes (--flag-filter-data True) and 400 control gene sets for score normalization (--n-ctrl 400). Downstream analyses were performed with the perform-downstream function, including group-level enrichment analysis across Seurat clusters (resolution 0.425) and gene-level association analysis, to identify cell populations and genes driving the disease relevance signal. Scores and statistics were imported from resulting .csv files into R and visualized as FeaturePlots() in Seurat [131,132].

### Immunofluorescence

D8 neuroectodermal spheres (NECs) were collected at day 8 of the differentiation protocol and washed 2x with PBS (D8537, Sigma-Aldrich). Afterwards, the NECs were fixed in 4% PFA (15710, Electron Microscopy Sciences) diluted in PBS for 1 hour at room temperature followed by two washing steps in PBS. NECs were either stored at 4°C in PBS until further usage or directly used to proceed with the IF staining. Therefore, the NECs were permeabilized in PBS supplemented with 0.5% Triton X-100 (9036-19-5, Merck) for 90 minutes at room temperature followed by an 1 hour blocking step in 1 ml blocking solution (0.5% Triton X-100, 5% BSA (8076.2, Roth), PBS) at room temperature. For stainings including SOX10 the blocking solution was additionally supplemented with 5% donkey serum (ab7475, abcam). Incubation with primary antibody dilutions (anti-SOX10 5 µg/mL, AF2864 R&D systems, anti-TFAP2A 0.4 µg/mL, sc-12726, Santa Cruz Biotechnology, lnc.) in blocking solution was performed over night at 4°C. The next day, the NECs were washed 3x for 15 minutes with PBS before adding the secondary antibodies (0.5 µg/mL, donkey anti-mouse IgG H&L 594 ab150108 abcam, donkey anti-goat IgG H&L 488 ab150129 abcam) in blocking solution. Incubation with the secondary antibody was performed for at least 2 hours at room temperature. The cells were then washed once with PBS for 10 minutes, once with PBS + DAPI (1µg/ml, 62248 Thermo Fisher) for 20 minutes and finally again with PBS for 10 minutes. NECs were mounted on glass slides with RapiClear (RC149001, SunJin Lab) and stored at 4°C until imaging.

For SOX10 immunofluorescence staining of attached human CNCCs at day 11 of differentiation, cells were fixed as described above. Permeabilization was, however, performed for 10 minutes at room temperature, and blocking in PBS supplemented with 5% donkey serum (without BSA or additional Triton X-100) for 30 minutes at room temperature. Primary antibody incubation (anti-SOX10, AF2864, R&D Systems; 1.25 µg/mL), secondary antibody incubation (donkey anti-goat IgG H&L 488, 1:1000), and DAPI staining were performed as above, except that all wash steps were carried out 3x for 5 minutes with PBS + 0.05% Tween-20, the secondary antibody was incubated for 30 minutes at 37°C, and DAPI was applied for 1 minute. Cells were mounted with Fluoromount-G.

Imaging of NECs was performed at the AX confocal microscope at the Nikon imaging center at Heidelberg University using a 25x silicone oil objective. Attached human CNCCs at day 11 of the differentiation were imaged using a Leica DMI4000 B fluorescent microscope with a 63x oil objective. Subsequent analysis was done using ImageJ2 (Version 2.14.0/1.54p).

### Data visualization

Relevant data visualization approaches were already described together with the underlying analysis above. The colour scheme used was viridis [137]. Multiple plots were assembled with patchwork [138].

## Supplementary Figures

**Supplementary Figure 1.**
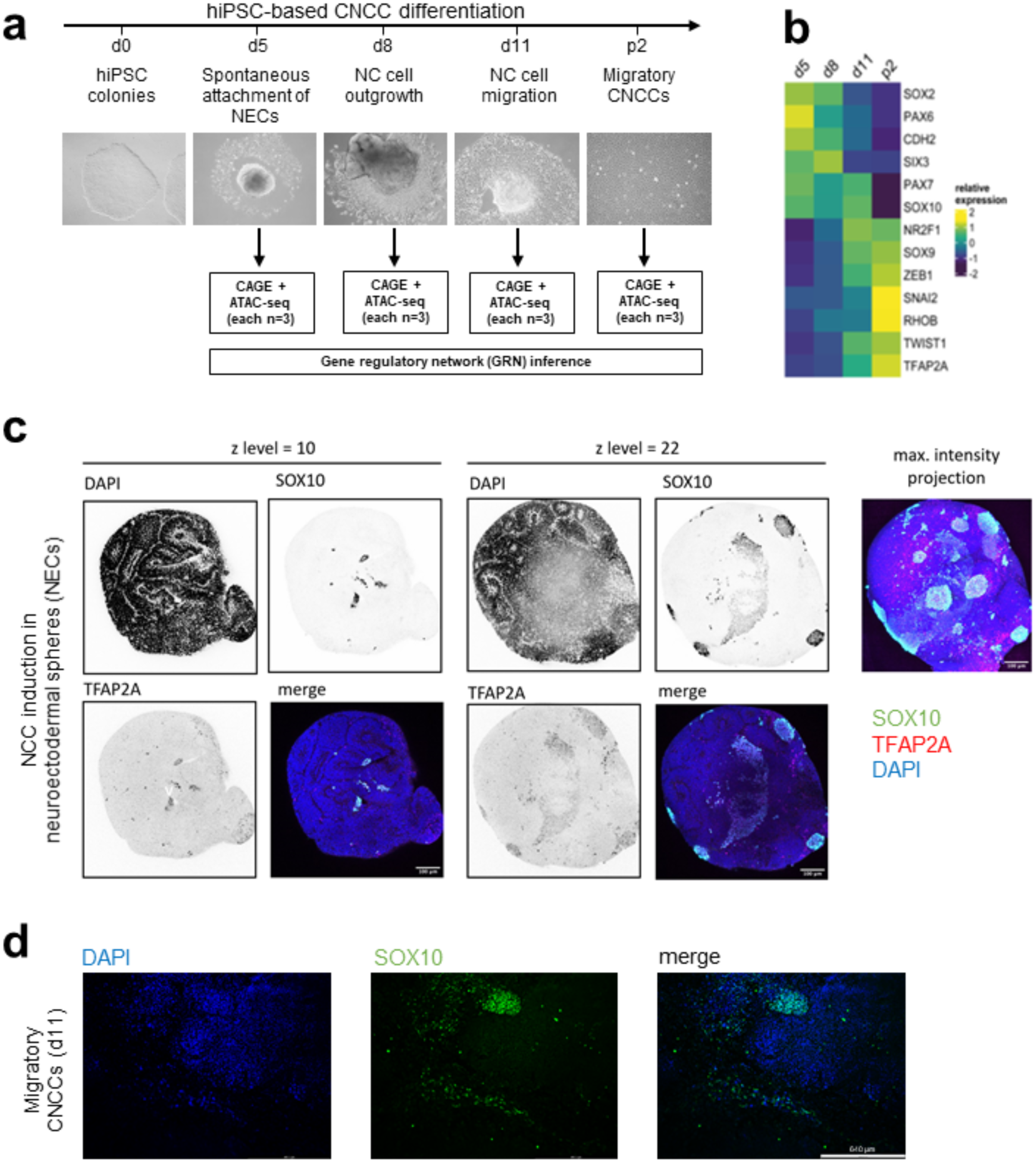
Data overview. **a** Schematic of the *in vitro* differentiation protocol of hiPSCs to human CNCCs with selected timepoints and representative brightfield microscopy images. For the indicated days, paired CAGE and ATAC-seq data (each n=3) was used to infer gene regulatory networks. **b** Expression of marker genes in the time-course CAGE data confirms the neuroectodermal stage at d5, NC induction and specification (d5-d11) and acquisition of a migratory CNCC identity (d11-p2). **c** Co-immunofluorescence staining of d8 NECs with SOX10 and TFAP2A in 25x magnification and nuclear stain using DAPI. Two different z-levels of the NEC are shown alongside a maximum projection of the full z-stack. **d** SOX10 staining of d11 CNCCs indicates a mixture of SOX10-positive and negative migratory cells.

**Supplementary Figure 2.**
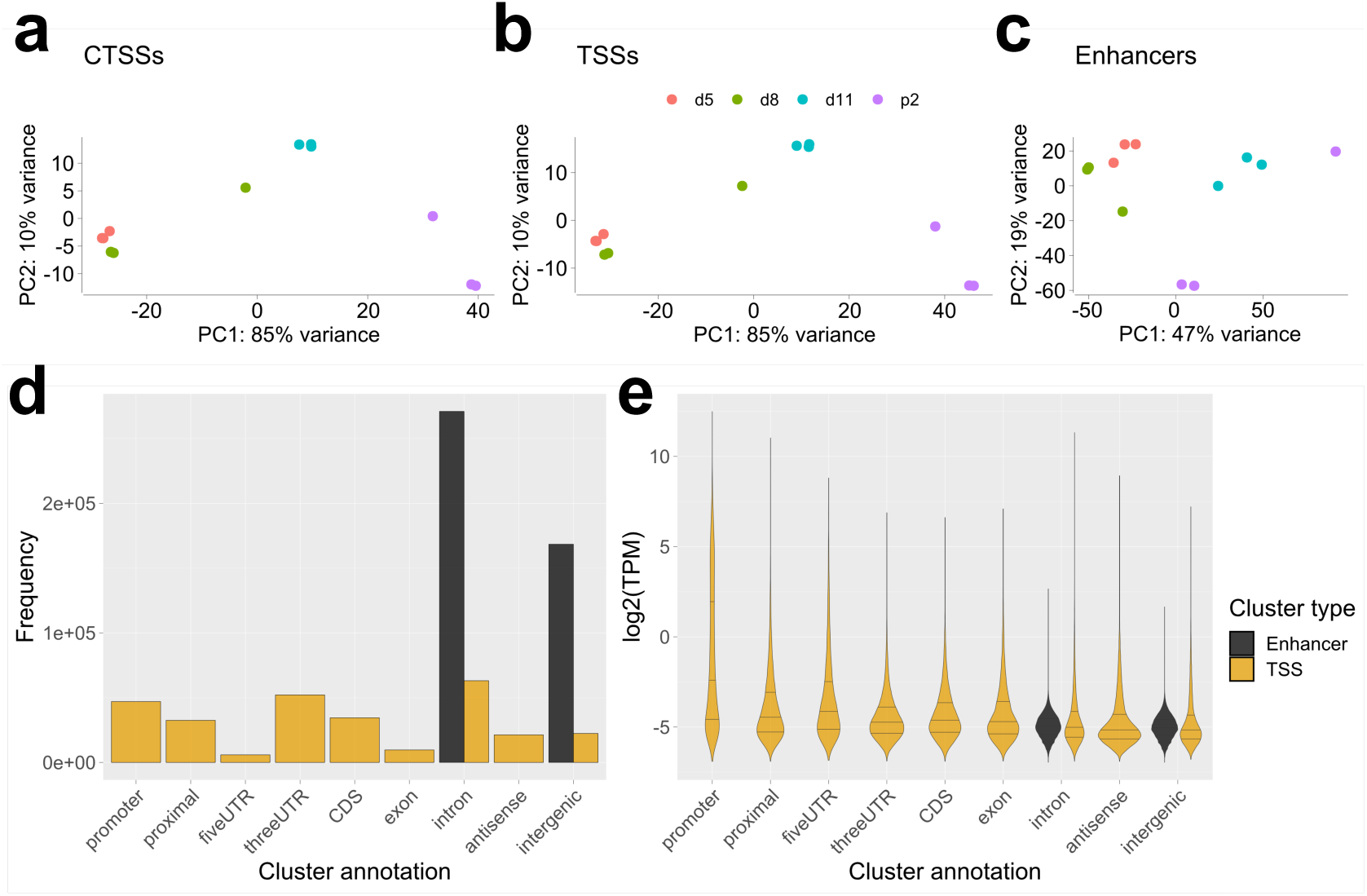
CAGE-seq preprocessing. **a** Principal component analysis (PCA) calculated from all CAGE-tags (CTSSs) **b** PCA calculated from all unidirectional clusters of CTSSs, i.e. putative transcriptional start sites (TSSs) **c** PCA calculated from all bidirectional clusters of CTSSs (putative enhancers). **d** Distribution of TSS and enhancers signals from CAGE data across genomic features. Notably, enhancer calling required them to map only to intronic or intergenic regions. **e** Expression levels of TSSs and enhancers across genomic features.

**Supplementary Figure 3.**
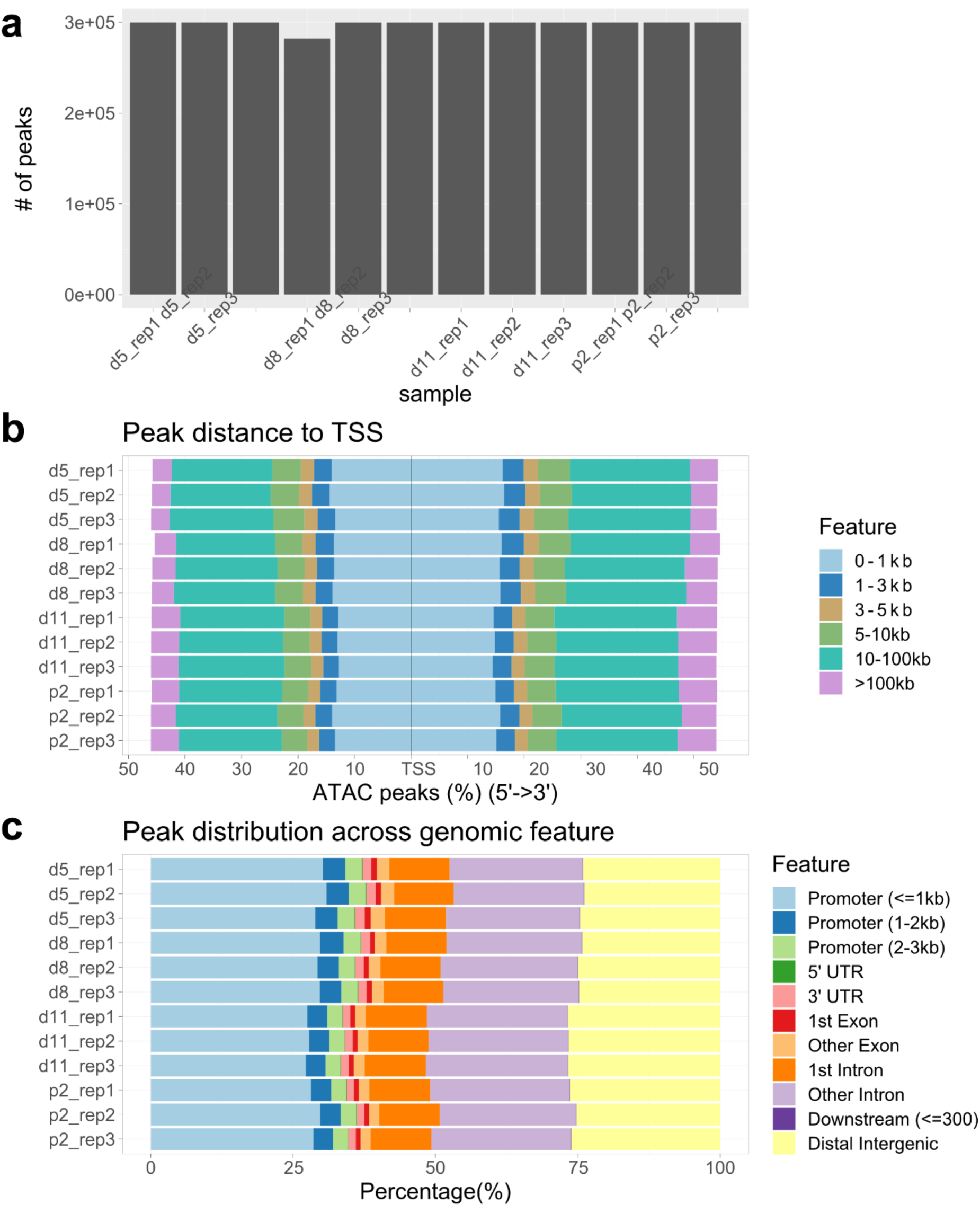
ATAC-seq quality control. **a** Number of identified peaks per sample **b** Distribution of ATAC-seq peaks around annotated transcriptional start sites (TSS). **c** Distribution of ATAC-seq peaks across genomic features.

**Supplementary Figure 5.**
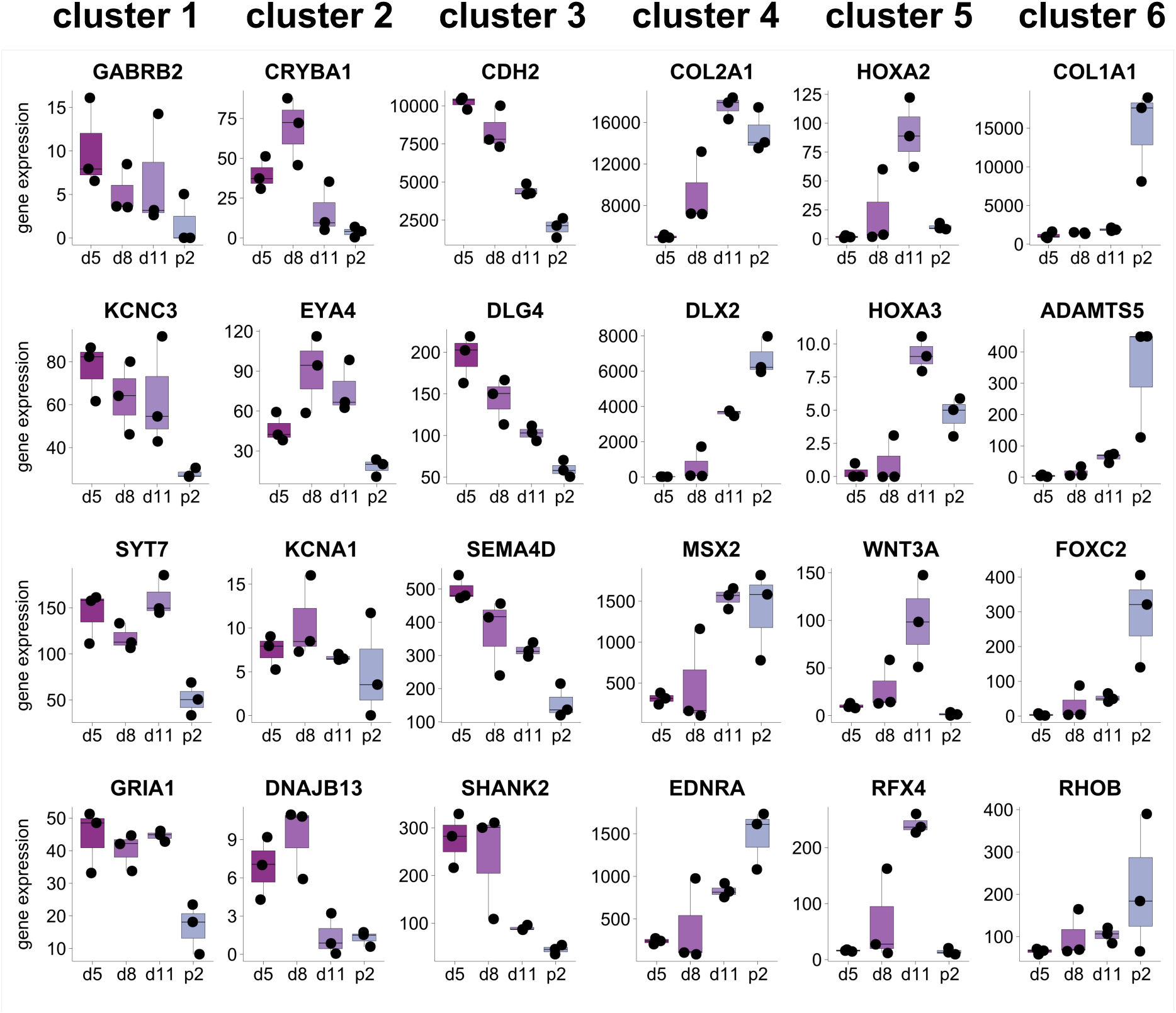
Example genes in clusters of dynamically expressed genes representative of the enriched processes. Genes expression from CAGE data for representative genes per cluster associated with the respectively enriched GO terms per cluster (Figure 1d).

**Supplementary Figure 6.**
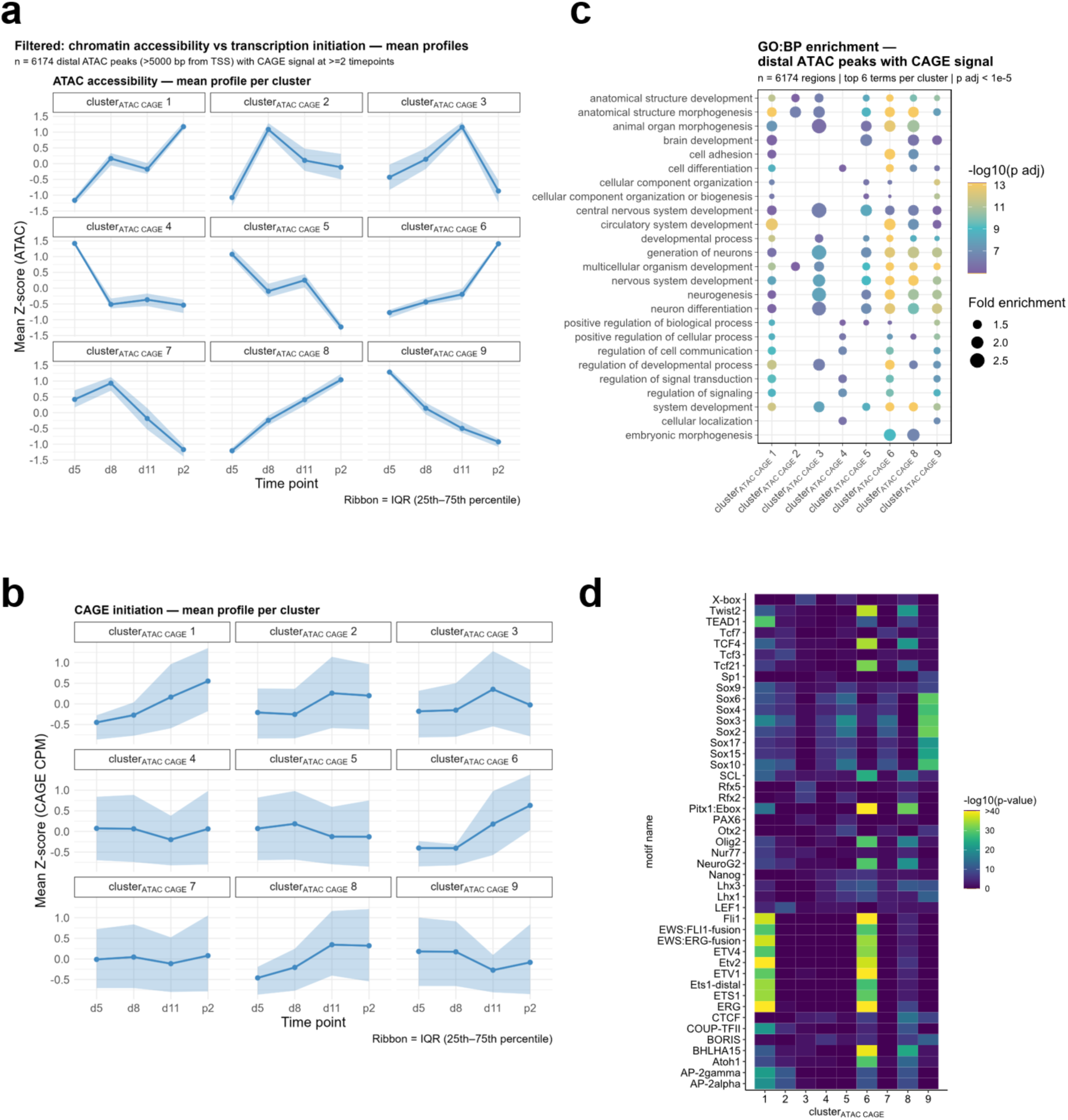
ATAC and CAGE data integration for identification of putatively active enhancers. **a** Regions from clustered ATAC-seq data were subset to distal regions (5000 bp from TSS) and those with CAGE signal in at least two time points. Average accessibility profiles with interquartile range are shown. **b** CAGE signal was summed for each accessible region and the average dynamics of CAGE counts per million (CPM) are shown for the regions corresponding to those in a). **c** Cluster-wise GREAT analysis shows processes governed by these putative enhancers. **d** Cluster-wise HOMER transcription factor motif enrichment analysis.

**Supplementary Figure 7.**
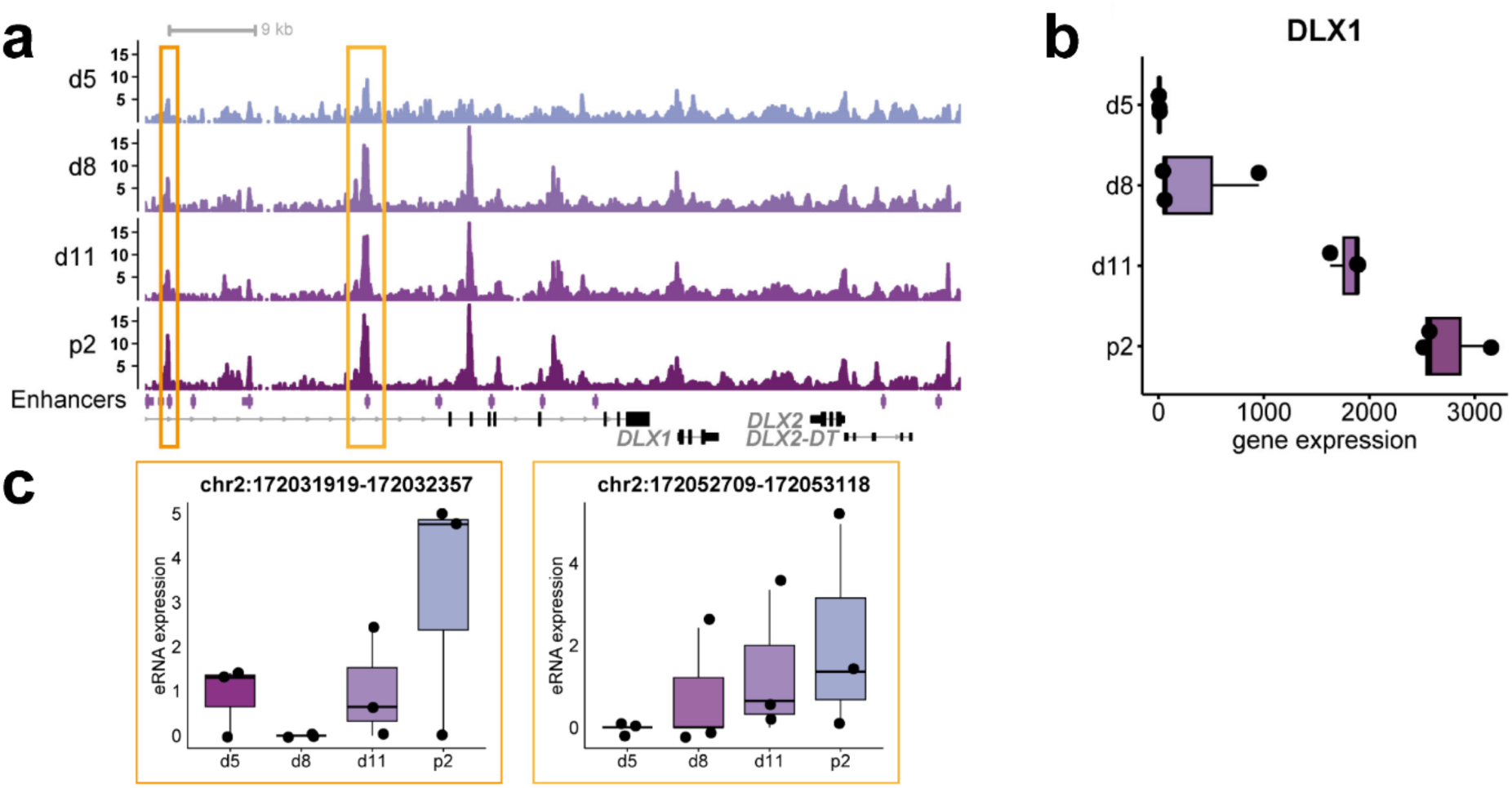
Correlation of gene and eRNA expression and chromatin accessibility changes supports RE-TG links. **a** Chromatin accessibility dynamics at the *DLX1*/*DLX2* locus during human CNCC differentiation. CAGE-identified enhancers are shown as a track underneath the ATAC-seq tracks. **b** *DLX1* expression dynamics during the differentiation. **c** eRNA expression dynamics at the two enhancer regions indicated by the orange and gold rectangles.

**Supplementary Figure 8.**
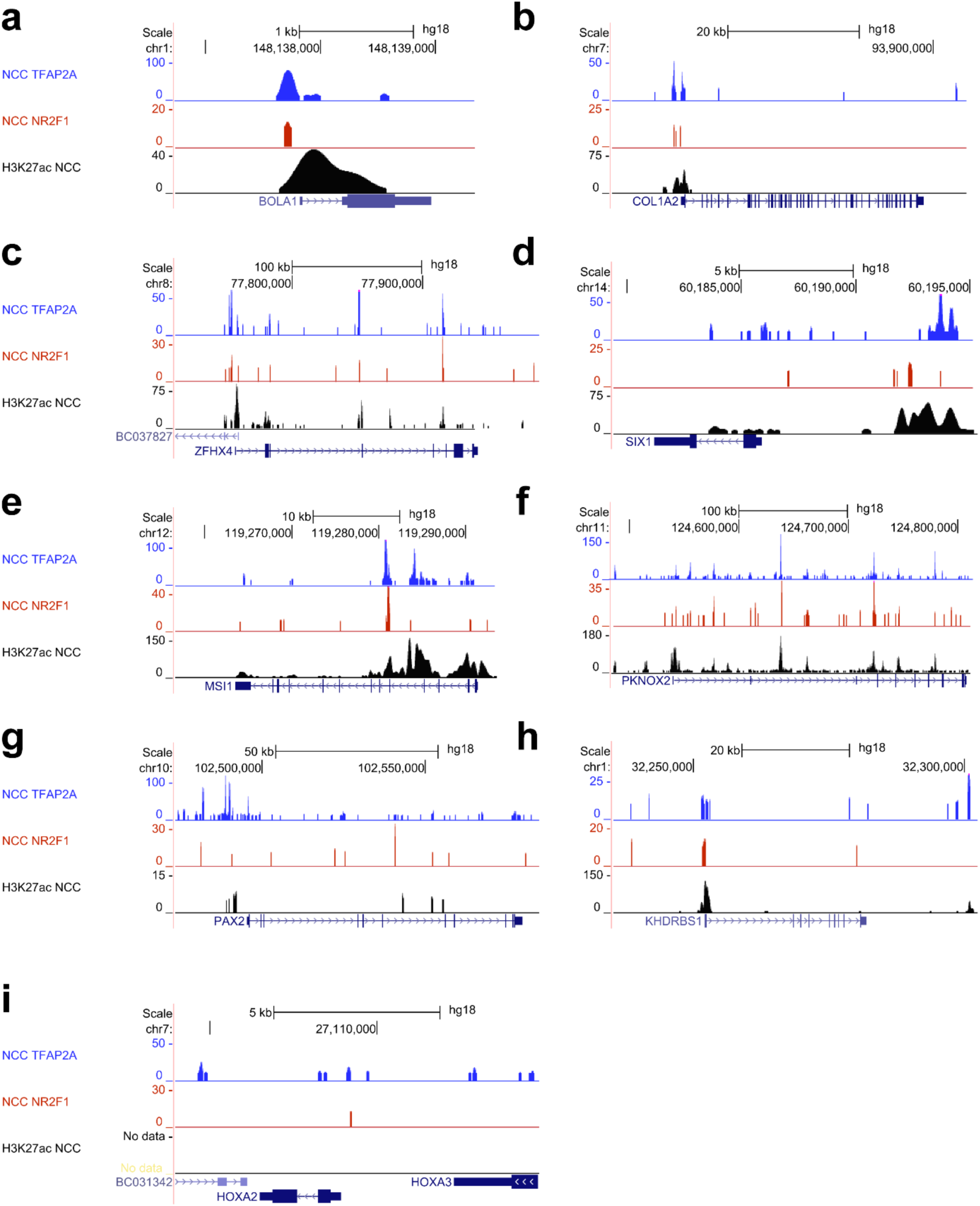
Published NR2F1 and TFAP2A ChIP-seq data from d11 confirms PECA2 prediction of NR2F1 and TFAP2A target genes on d11 and p2 for high trans-regulation scores. Note that LPAR4 in **(j)** is not a predicted target of NR2F1.

**Supplementary Figure 9.**
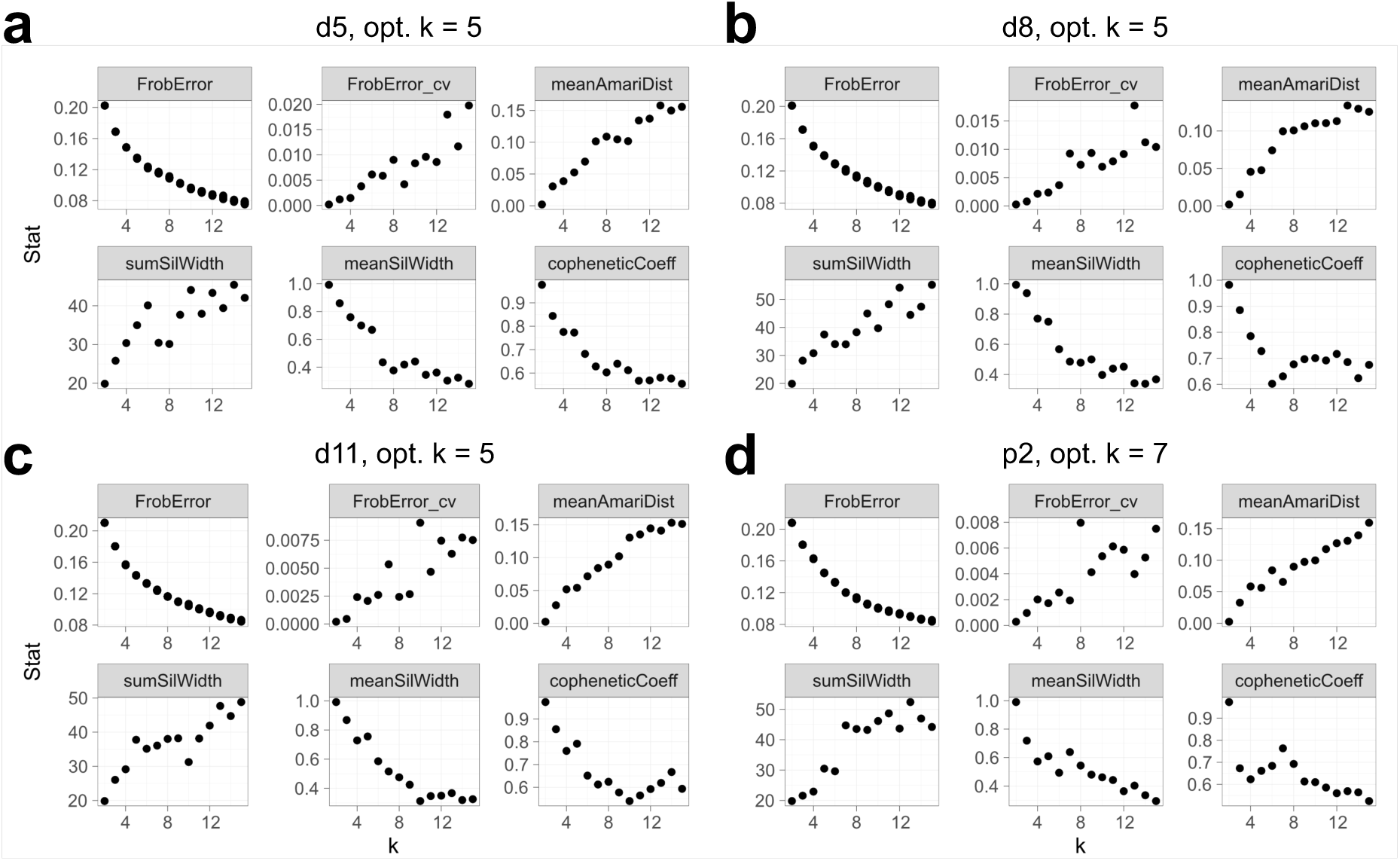
Determination of optimal factorization rank for NMF per time point. Six measures to determine the optimal factorization rank for NMF for d5 (**a**), d8 (**b**), d11 (**c**) and p2 (**d**).

**Supplementary Figure 10.**
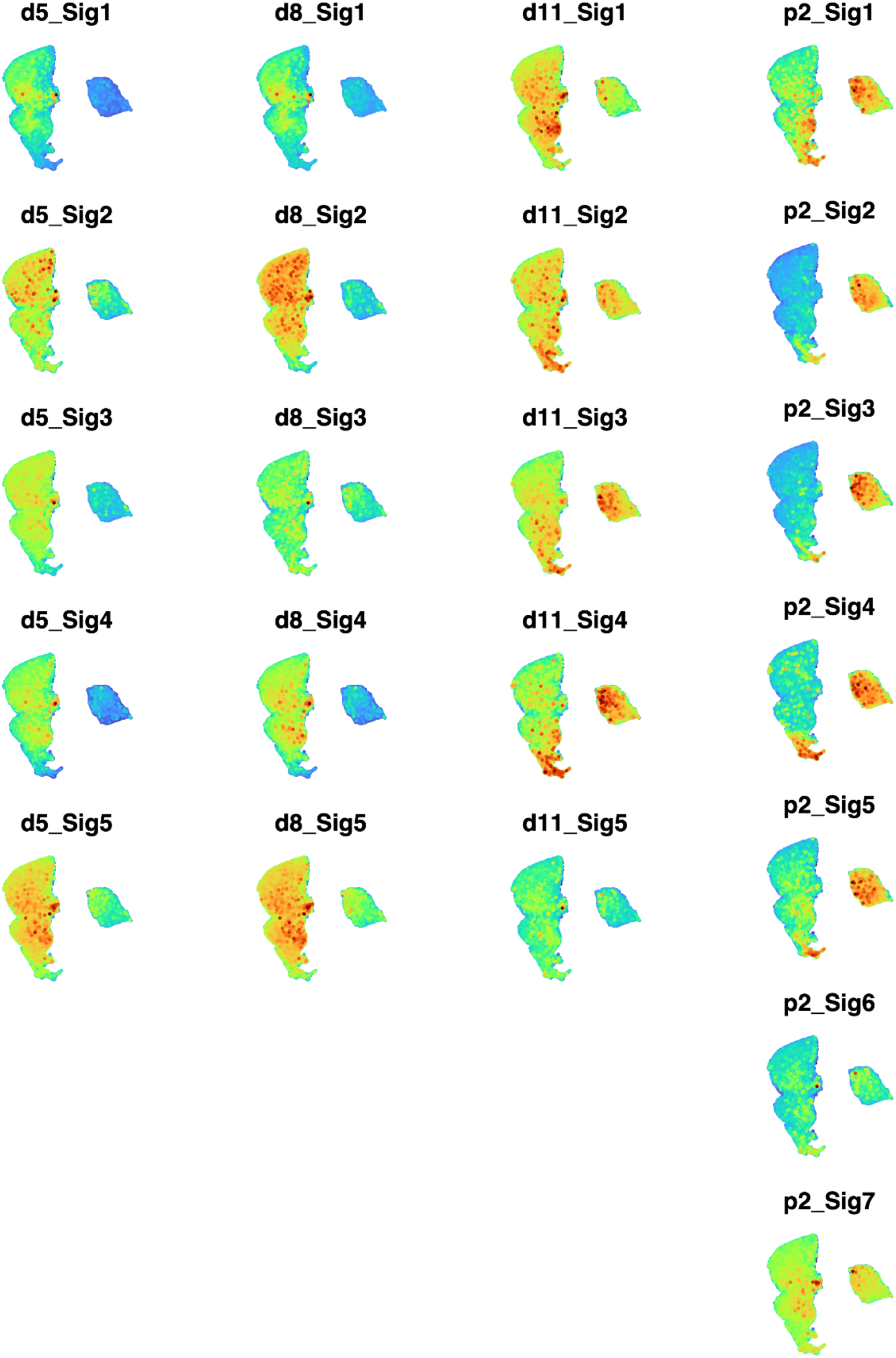
Scoring of modules obtained from NMF of trans-regulation score matrices per day. NMF-identified core regulatory modules were scored in the scRNA-seq data using AUCell.

**Supplementary Figure 11.**
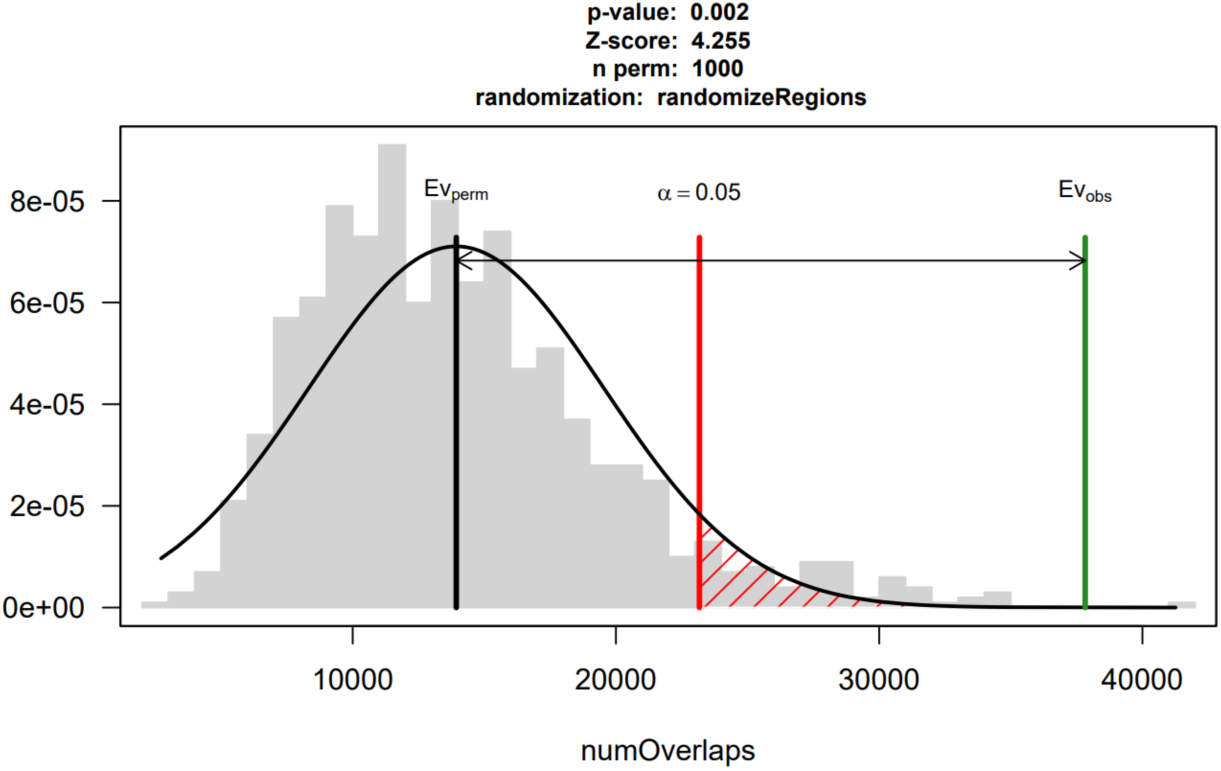
nsCL/P risk regions significantly overlap with REs in the obtained GRNs governing human CNCC differentiation. Permutation testing shows significant enrichment of nsCL/P risk regions among regulatory elements in our GRNs.

**Supplementary Figure 12.**
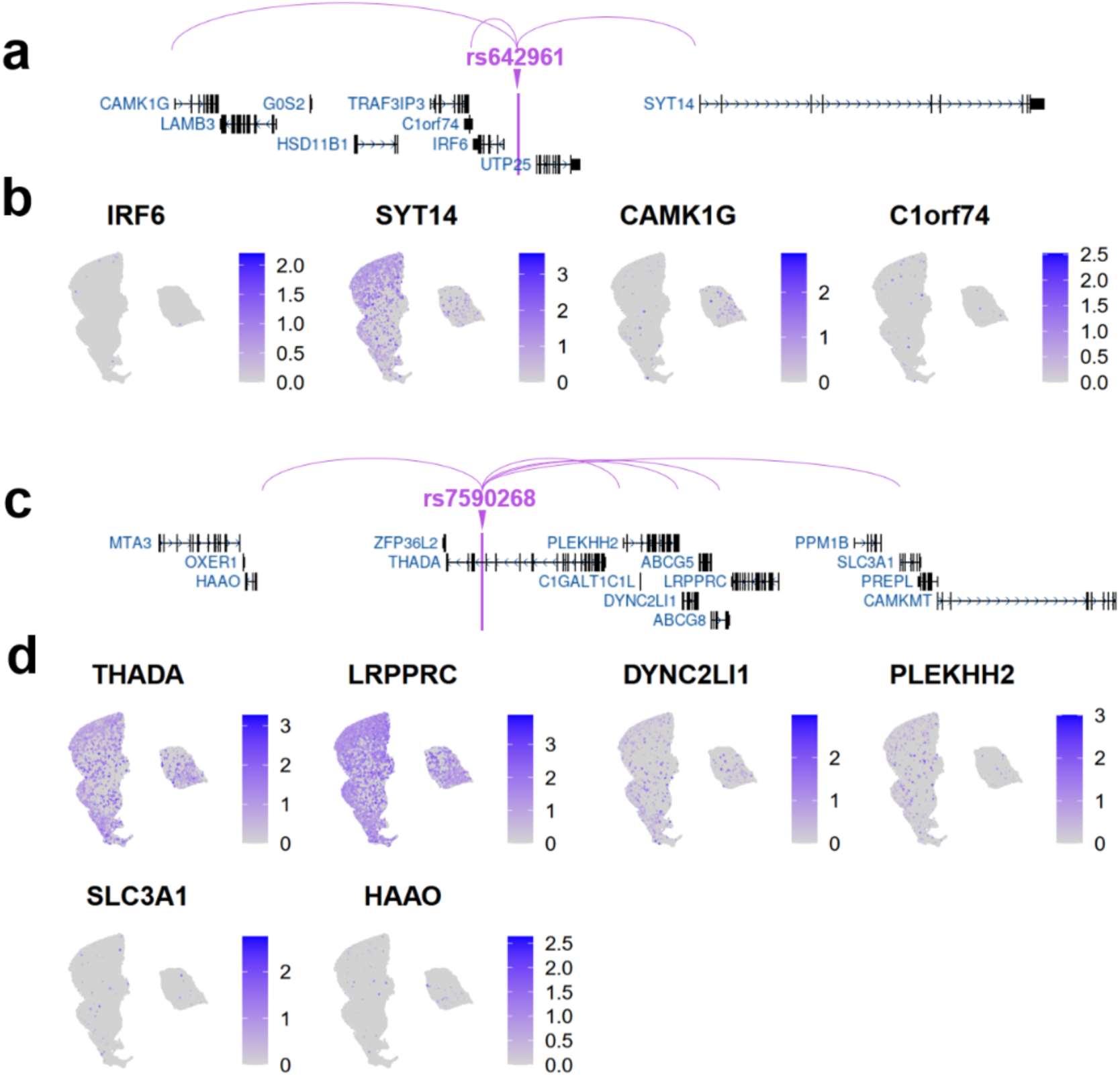
Example nsCL/P-associated SNP-TG links according to the PECA2 predictions with expression of predicted targets and locus genes underneath. A schematic that connects the lead variant for each risk locus with the predicted TGs and expression dynamics underneath for the *IRF6* locus (**a**, **b**) and the *THADA* locus (**c**,**d**).

